# A body detection inversion effect revealed by a large-scale inattentional blindness experiment

**DOI:** 10.1101/2024.10.18.619010

**Authors:** Marco Gandolfo, Marius V. Peelen

**Affiliations:** Donders Institute for Brain, Cognition, and Behaviour, Radboud University, Nijmegen, The Netherlands

**Keywords:** attention, expectation, social cognition, body perception, configural processing, inversion effects, inattentional blindness

## Abstract

As a social species, humans preferentially attend to the faces and bodies of other people. Previous research revealed specialized cognitive mechanisms for processing human faces and bodies. For example, upright person silhouettes are more readily found than inverted silhouettes in visual search tasks. It is unclear, however, whether these findings reflect a top-down attentional bias to social stimuli or bottom-up sensitivity to visual cues signaling the presence of other people. Here, we tested whether the upright human form is preferentially detected in the absence of attention. To rule out influences of top-down attention and expectation, we conducted a large-scale single-trial inattentional blindness experiment on a diverse sample of naive participants (N=13.539). While participants were engaged in judging the length of a cross at fixation, we briefly presented an unexpected silhouette of a person or a plant next to the cross. Subsequently, we asked whether participants noticed anything other than the cross. Results showed that silhouettes of people were more often noticed than silhouettes of plants. Crucially, upright person silhouettes were also more often detected than inverted person silhouettes, despite these stimuli being identical in their low-level visual features. These results were replicated in a second experiment involving headless person silhouettes. Finally, capitalizing on the exceptionally large and diverse sample, further analyses revealed strong detection differences across age and gender. These results indicate that the visual system is tuned to the form of the upright human body, allowing for the quick detection of other people even in the absence of attention.

## 1. Introduction

Object perception starts with detecting the presence of an object (is there something there?), followed by the recognition of that object (what object is there?). A long-standing question in vision science is whether these processes rely on common or distinct mechanisms (de la Rosa et al., 2011; Grill-Spector & Kanwisher, 2005; M. L. Mack et al., 2008; Nakayama et al., 1995; Rosch et al., 1976). Evidence shows that familiar inanimate objects presented in their canonical orientation can be recognized as quickly as they can be detected. For example, as soon as the presence of a plant is detected, people can also recognize that object as a plant (Grill-Spector & Kanwisher, 2005). However, other research demonstrates that detection and recognition mechanisms can be dissociated: when an object appears inverted, or degraded, recognition performance is slowed down, while detection performance remains unaffected (M. L. Mack et al., 2008). As such, it is likely that detection is primarily determined by low-level visual features (e.g., contrast, texture) that are not disrupted by inversion.

The effect of inversion on recognition performance famously differs between object categories (Yin, 1969), with face and body recognition disproportionally affected by inversion relative to other object categories (Carey et al., 1992; Farah et al., 1995; Reed et al., 2003; R. A. Robbins & Coltheart, 2012b; Yin, 1969). These findings have been interpreted as reflecting configural processing of faces and bodies and have been used to argue for specialized mechanisms underlying face and body recognition (Farah et al., 1998).

Interestingly, there are two lines of evidence suggesting that *detection* mechanisms may also be specialized for faces and bodies. The first comes from studies using continuous flash suppression (CFS), a method that renders an object presented to one eye temporarily invisible by simultaneously presenting a high contrast mask to the other eye (Jiang et al., 2007; Tsuchiya & Koch, 2005). Upright faces and bodies “break” through the mask and are detected more quickly than inverted faces and bodies (Jiang et al., 2007; Stein et al., 2012). The detection inversion effect is specific to faces and bodies (Jiang et al., 2007; Stein et al., 2012). Because the low-level features between upright and inverted stimuli are identical, the detection inversion effect for person stimuli raises the intriguing possibility that bottom-up detection mechanisms may already be tuned to human face and body configuration.

However, an alternative explanation for these CFS findings is that they reflect the modulatory effect of attention, known to affect interocular suppression (Paffen & Alais, 2011; Stein & Peelen, 2015). Participants in these experiments performed hundreds of trials, and were fully aware of the stimulus set (i.e., the faces and/or bodies) presented in the experiment. Studies have shown that attention can be selectively directed to (upright) faces and bodies, and that this modulates early stages of visual processing, similar to attention to low-level features (Störmer et al., 2019; Thorat & Peelen, 2022). Therefore, the detection inversion effect for person stimuli may be mediated by a top-down process of increased anticipation for upright depictions of the human figure, rather than reflect bottom-up sensitivity. Furthermore, it is possible that the previously reported detection inversion effect is specific to the CFS method, in which objects are suppressed for several seconds, potentially allowing for extensive processing before the detection response (Stein, 2019; Stein & Peelen, 2021).

The second line of evidence for specialized detection of person stimuli comes from inattentional blindness studies, showing that faces and bodies are more likely to be noticed than other object categories when attention is directed elsewhere and no stimulus is expected to appear (Devue et al., 2009; Downing et al., 2004; A. Mack & Rock, 1998). In these studies, participants judged the length of one arm of a visually-presented cross. In a critical trial, unbeknownst to the participants, an unexpected stimulus (e.g., a person silhouette) appeared nearby the cross. Directly afterwards, participants were asked whether they noticed a stimulus. Participants were more likely to notice the presence of a face or a person silhouette than the presence of other object silhouettes (e.g., tools). Importantly, in this task (A. Mack & Rock, 1998), unlike in the CFS paradigm, the stimuli are presented binocularly and participants have no knowledge of, or expectation for, any stimulus to appear. These findings therefore suggest that detection mechanisms may be sensitive to person stimuli even in the absence of top-down attention and expectation.

However, these results are not straightforward to interpret because detection rates were compared across object categories (e.g., person vs car; Downing et al., 2004) without an inverted version of these stimuli. Before being affected by an object’s category, detection sensitivity is primarily sensitive to low-level stimulus features such as contrast, luminance, and spatial frequency. Different object categories may show different detection rates based on such features. Indeed, a recent metanalysis on inattentional blindness showed that visual features are stronger moderators of detection rates than higher-level stimulus properties (Hutchinson et al., 2022). Furthermore, due to the single-trial nature of this paradigm, it has been challenging to obtain data with enough statistical power as this would require testing hundreds of participants (for a face inversion experiment in a small sample, see Devue et al., 2009). As such, it remains an open question whether detection mechanisms are sensitive to the configuration of person stimuli.

Here, we addressed this question by comparing detection performance for upright and inverted body silhouettes. We made use of large-scale data collection outside the university setting to provide a definitive answer to whether human bodies are preferentially detected in the absence of expectations, and while controlling for low-level visual feature differences via inversion. Mack and Rock’s (1998) inattentional blindness task (see **Figure 1**) was presented to unsupervised visitors of a museum (MuZIEum, Nijmegen, The Netherlands) in the context of an educational demonstration about attention. Testing this task in a museum allowed for measuring detection rates in a fully naïve population, which may not be achieved through online crowdsourcing platforms (Stewart et al., 2017). It also allowed for testing an exceptionally large and diverse sample (N = 13539; age range: 11-85); to our knowledge the largest sample ever employed in an inattentional blindness experiment. Participants judged the length of a cross briefly presented on the screen. Unbeknownst to the participants, in critical trials an upright or inverted body or plant silhouette appeared nearby the cross. Detection rates of the stimuli were measured as a function of Category (person/plant) and Orientation (upright/inverted). If the visual system contains specialized detection mechanisms for upright person stimuli, then upright person silhouettes should be noticed more often than their inverted, visually matched counterparts, even when these stimuli are not expected, and not task relevant. Further, inversion effects for detection should be specific to bodies, and not reliably observed with a control object category (i.e., plant stimuli) in which detection likely relies only on low-level features not affected by inversion (Mack et al. 2008).

**Figure 1.**
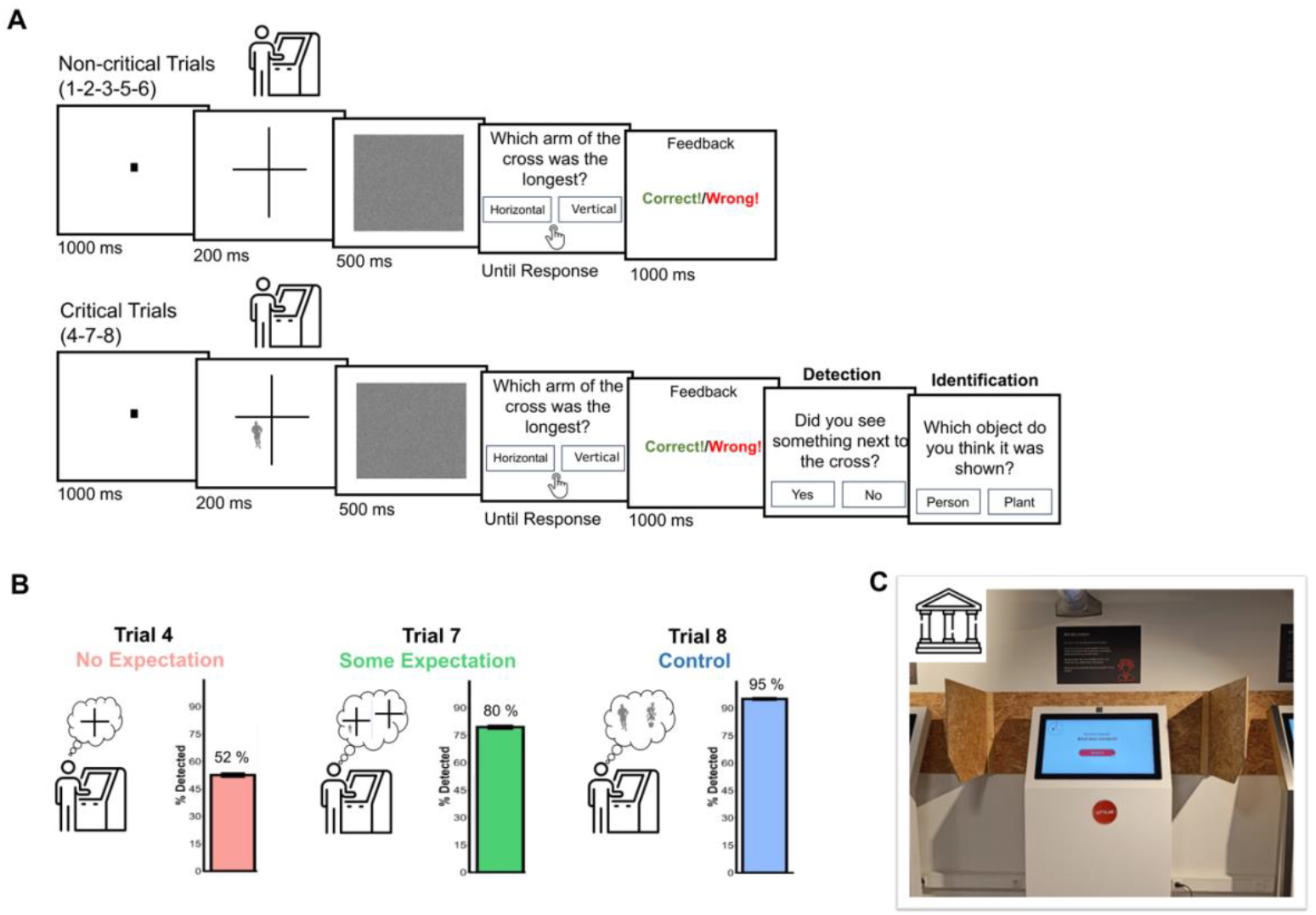
**A.** Schematic representation of the trial procedure of the Mack and Rock’s cross task. The top row shows the procedure for the non-critical trials (1-2-3-5-6). The bottom row shows the procedure for the critical trials (Trial 4-Trial 7-Trial 8), in which the stimulus appeared inside the cross and participants were asked a detection and an identification question after the feedback on the cross-task answer. **B**. Schematic representation of the three critical trials. In Trial 4 participants had no expectation and had not yet been asked about, or presented with, a stimulus inside the cross. In Trial 7, participants were not alerted that a stimulus would appear in the cross but formed some expectations about this possibility based on the detection and identification questions asked in Trial 4. In Trial 8, participants were explicitly alerted that something would appear inside the cross before the trial started. Alongside the representation of each trial, the bar plot shows the average detection rates across stimuli. In trial 4, the stimulus was detected 52% of the times, meaning that 48% of our sample (N = 6480 people) was inattentionally blind to the stimulus in this trial. The detection rates reliably increased in trial 7 (80%) and increased even further in the control trial, trial 8 (95%). **C**. Example of the museum setup. The experiment ran on a touch-screen kiosk in the museum exhibition room. Participants were museum visitors who performed the experiment on a voluntary basis by approaching the kiosk. Above the screen there was a brief description of the experiment and information about its duration.

In addition, we capitalize on the exceptionally large and diverse sample to assess age and gender differences in inattentional blindness. First, we assessed whether these demographic variables modulated the detection rates depending on category and orientation of the silhouettes appearing inside the cross. Second, as recommended by a recent metanalysis of inattentional blindness (Hutchinson et al., 2022), we report how these differences influenced detection rates across stimuli, since demographic information has often been ignored in previous work (Hutchinson et al., 2022), and has led to mixed evidence in smaller samples (Graham & Burke, 2011; Hannon & Richards, 2010; Kreitz, Furley, et al., 2015; Kreitz, Schnuerch, et al., 2015; Richards et al., 2010; Stothart et al., 2015; B. B. Velichkovsky & Popova, 2021).

## 2. Methods and Materials

### 2.1 Participants

An essential requirement for inattentional blindness studies is that subjects are naïve as to the purpose of the experiment. Indeed, university students and online platforms workers (e.g., mTurk, Prolific) are often highly experienced in performing experiments and may thus not be entirely naïve. In their seminal work, Mack and Rock (1998) refer to an “insatiable need” of naïve subjects. Similarly, to ensure a large, diverse, and naïve sample we ran our study on visitors of a local museum (muZIEum, Nijmegen, The Netherlands), within the context of a demonstration on attention.

Participants freely approached the experiment in the museum hall. The final sample considered for the analyses includes data from N = 13539 participants (N = 8416 identifying as woman, and N = 5123 identifying as man; mean age = 33.0 ± 16.5; age range: 11-85 years). Details about participants’ inclusion and exclusion criteria are provided in the Data analysis section. The data were collected between February 2022 and the end of September 2024. All participants gave consent for their data being used for scientific research. The procedures obtained approval from the Ethics Committee of the Faculty of Social Sciences at Radboud University (ECSW-2021-076).

### 2.2 Stimuli and apparatus

The stimuli were selected from the stimulus pool of Stein and Peelen (2012). Specifically, they were 40 person silhouettes and 40 silhouettes of house plants with luminance of 50% (middle gray RGB – 128, 128, 128). To increase within-category variability, the body silhouettes depicted various poses, such as standing, running, dancing, exercising, sitting, jumping, and they were seen from different viewing angles. Similarly, the plants varied for their species (e.g. Ficus, banana plant, papyrus). To add further stimulus variability, we also generated a horizontally flipped version for each of these 80 images. Finally, for each image we generated an equivalent number of inverted stimuli by rotating each image by 180°.

The cross was generated by intersecting a vertical and a horizontal black line. The length of the lines randomly varied between 400 and 480 pixels in intervals of 20 pixels. The largest difference between the two lines on the screen was of ∼3cm. The mask consisted of Gaussian white noise.

The experiment ran on a 32-inches kiosk touchscreen (1920 x 1080 pixels) placed in the hall of the muZIEum, a museum focused on increasing awareness of visual impairments. The experiment was part of the project “Donders Citylab” (see **Figure 1C** for an example of the setup). The experiment was coded in JavaScript using jsPsych (de Leeuw, 2015) and its psychophysics toolbox (Kuroki, 2021). The data were hosted on Pavlovia.org servers.

### 2.3 Procedure

The experiment ran on one of three kiosk screens in the museum hall (see **Figure 1C**). Participants approached it freely during their visit. A welcome screen showed the duration of the experiment (∼2.5 minutes, including instructions and debriefing) and a button to start it. When participants started the experiment, they then selected their preferred language (Dutch – 85% of the participants, or English – 15% of the participants), indicated their age on a touch-sensitive slider, and (if they wanted) their gender. They reported their gender with a multiple-choice question with the following options: man, woman, other, prefer not to say. Crucially, in a following multiple-choice question we also asked whether participants knew about the topic of the demonstration and/or participated in the experiment before. This was to filter out datafiles including participants who may were not naïve to the inattentional blindness phenomenon, or who wanted to repeat the demonstration again, after having read the debriefing.

#### Mack & Rock’s inattentional blindness task

Written task instructions were presented in the context of a family-friendly explanation of the cocktail party phenomenon, that is, our ability to focus on certain information very effectively. Participants were then informed that the experiment aimed to measure the ability to focus on a task, and that this ability varies between individuals. The cross task was then explained with a gif of the trial procedure running at the same speed of the task.

Each trial started with a 1 second fixation dot followed by a cross (200 milliseconds), and by the mask (500 milliseconds). After the mask, a response screen appeared asking “which arm of the cross do you think is the longest?”. Participants could choose between one out of two buttons on the touch screen: “Horizontal” and “Vertical”. After each response, they were given feedback about their performance.

In the critical trials (Trial 4, Trial 7, and Trial 8), right after the cross response, two further questions were asked on a button response screen: 1) A detection question - “Did you see something next to the cross?” (yes or no); and 2) An identification question - “Which object do you think was shown?” (person or plant). Finally, in the last trial (Trial 8) participants were instructed to identify the object inside the cross before the cross appeared (see **Figure 1**).

The experiment ended with a more detailed explanation of the inattentional blindness phenomenon, explaining the cost of selectively paying attention to a stimulus. Additionally, participants also had the option to be fully debriefed about our specific hypotheses (i.e., that bodies are detected more often than other objects) by scanning a QR code with a link to an outreach article on inattentional blindness (written in Dutch and English, https://blog.donders.ru.nl/?p=13893&lang=en).

### Design

The experiment was a between-participants design similar to Downing et al. (2004). Each participant performed 8 trials (see **Figure 1AB**). In the critical Trial 4, Trial 7, and Trial 8 a stimulus appeared in one random quadrant of the task-relevant cross (**Figure 1B**). Trial 4 represents the purest measure of inattentional blindness, because participants should have no expectation of the stimulus appearing inside the cross. Trial 7 is often called “divided attention trial” (Mack & Rock, 1998), in that participants are already aware of the possible presentation of a stimulus nearby the cross, though not explicitly alerted that it would appear on that specific trial, or again in the experiment. Indeed, in this trial, as opposed to trial 4, participants have at least *some* expectation of the possible appearance of a stimulus on the screen. Finally, trial 8 is often termed as the “full attention trial”: here, participants are explicitly told to ignore the cross and identify the object that appears within it. This serves as a control to ensure that the stimuli can be identified when they are the sole focus of attention.

Each participant was randomly assigned to one out of four types of stimuli (person upright/person inverted/plant upright/plant inverted) in each of the three critical trials (Trial 4, Trial 7, and Trial 8). Importantly, none of these four stimulus-by-orientation combinations were repeated within the same participant, nor was the same stimulus exemplar repeated within participant.

### 2.5 Analyses

First, we compared detection rates between the critical trials across category and orientation to validate the task on the large and diverse sample of unsupervised museum visitors. We then used logistic regression models to assess whether detection rates differed by category and orientation in both Trial 4 (No-Expectation) and Trial 7 (Some-Expectation). Details on model selection are available in the **Supplementary Material**. For the selected models, we report the model’s estimate values for each factor (exponentiated coefficients), the 95% confidence interval of these coefficients, and the standardized betas. In the presence of an interaction, we followed up with chi-squared tests contrasting detection rates of upright vs inverted within each object category. To allow inference on the absence of effects for chi-squared contrasts we performed Bayesian chi-squared tests using the function *contingencyTableBF* of the R package BayesFactor (Morey et al., 2024). Bayes factors (BF_10_) below 1 indicate that the null hypothesis is more likely, and values below 1/3 (0.33) are generally taken as reliable evidence in favor of the null hypothesis. Conversely, BF_10_ above 3 indicate evidence for the alternative hypothesis (Jeffreys, 1961; Lee & Wagenmakers, 2014; Ly et al., 2016). The same analyses were performed for the identification rates, separately for Trial 4 and Trial 7.

Finally, following previous studies assessing age and gender differences in inattentional blindness (Hannon and Richards, 2010; Kreitz, Schnuerch et al., 2015; Richards et al., 2010; Stothart et al., 2015; Graham and Burke, 2011; Kreitz et al., 2015; Velichkovsky et al., 2021), we ran logistic models to test the role of these individual differences on detection rates across all the other factors.

All the analyses were conducted using R (*R: core team*) version 4.4.1. The data were processed and visualized using the package *tidyverse* (Wickham et al., 2019) and *patchwork* (Pedersen, 2024). The exponentiated coefficients for the regressions were calculated using package *jtools* (Long, 2024) and the oddsratios on the chi-squared comparisons were calculated with package *epitools* (Aragon et al., 2020). The ANOVAs and t-tests were performed using package *bruceR* (Bao, 2024).

#### 2.5.1 Participant exclusions

For the analyses we considered the population aged between 11-85 years old (N = 18117 data files reporting age within this range). This was the largest plausible age range because people younger than 11 years old would not be likely to reach the height of the kiosk screen to efficiently perform the task and people above 85 may not faithfully indicate their age (indeed we noted an unlikely peak at 100 years old and very low overall numbers of datafiles above 85 – see **Figure S1** in the **Supplementary Material**). For the same reason, we also excluded participants who did not move the age slider from the default value of 50 (N = 763, 4.21% of the total number of datafiles). We observed an unexpected and unlikely peak of datafiles where this “default” age was indicated (See **Figure S1**). We then excluded participants who declared to know about the demonstration (i.e., the phenomenon of inattentional blindness) or declared to have participated before or wanted to repeat the experiment (N = 3069, 16.94% of the total no. of datafiles). After these exclusions we had N = 14285, and then considered outliers for performance in the cross task. We excluded participants who did not answer correctly in the cross task in at least 2 out of 8 trials (N = 174, 1.21%), and participants who across conditions had average response times in the cross task (RT) 2.5 SDs slower than the group mean (mean RT > 6.37 seconds, N = 166, 1.16%). To facilitate regressing out and to assess gender effects on a reasonably balanced sample we excluded datafiles in which the reported gender was “other” (i.e., neither man nor woman, N = 151, 1.06%) or that preferred to not report their gender (N = 253, 1.77%). For transparency, we made available all the data files in the **OSF repository** (https://osf.io/ahm7r/) and include the code that led to these exclusions.

To further ensure data quality, when comparing the detection rates for each category and orientation with logistic regression, we further excluded those participants who answered “No not detected” in the *detection* question of Trial 8 (i.e., the full attention trial; Mack and Rock, 1998; N = 675, 4.99% of the remaining datafiles). Indeed, in this trial the participants were explicitly told that a to be identified stimulus would appear inside the cross. The lack of detection of the stimulus inside the cross when this was fully expected was taken as a valuable attention check and follows the approach of previous studies of inattentional blindness (Cartwright-Finch & Lavie, 2007; Remington et al., 2014; for a metanalysis on the topic see Hutchinson et al., 2022). The pattern of results on detection rates did not change as a function of these exclusions. Separate analyses on detection rates and identification performance of Trial 8 are in the **Supplementary Material**, see **Figure S3**.

## 3. Results

### Inattentional blindness in museum visitors

While our recruiting method maximized the chance of finding naïve participants, it may reduce our chances to find inattentional blindness, due to the speeded presentation of the stimuli and the lack of experimenter supervision. Therefore, our first analysis goal was to validate Mack and Rock’s inattentional blindness task (1998) on unsupervised visitors of a museum.

The first validation measure was the comparison in terms of detection rates between the first critical trial (Trial 4) and what Mack and Rock defined as the control or full attention trial (Trial 8, see **Figure 1B**). The control trial is the last trial of the cross task. Here, before the trial starts, participants are instructed to not necessarily pay attention to the cross but to attend the object that appears inside it. As such, the control trial provides a measure of the perceptibility of the stimuli under the conditions of a brief, masked, parafoveal presentation but with full expectation towards the appearance of a stimulus. If the detection rates are significantly higher in Trial 8 than Trial 4, it means that the stimulus appearing in the cross is detected when it is expected and attended, in spite of the brief presentation of the display followed by the mask (200ms).

The difference between detection rates on the critical Trial 4 vs Trial 8 was significant (**Figure 1B**, *χ*^2^(1) = 6401.1, *p <* 0.001, v = 0.49, BF_10_ > 1000). In Trial 4, out of 13539 participants, 6480 participants (47.9%) failed to detect the stimulus while being engaged in the cross task. Conversely, in Trial 8, the stimulus was detected by 12864 participants (95.0%) and only 5% of the participants (n = 675) failed to detect it. The stimulus was also presented in Trial 7, also called the divided attention trial. In this trial, the detection rates for the stimulus are usually placed in between the first critical inattention trial (Trial 4) and the full attention trial (Trial 8). Indeed, in Trial 7, participants should have *some* expectation about a stimulus appearing nearby the cross, because they have been previously asked about it in Trial 4. Yet, they were not explicitly warned that a stimulus would be appearing again inside the cross as in Trial 8. As expected, in this trial, more participants (10779 out of 13539; 79.6%) than in Trial 4 detected the presence of the stimulus (*χ*^2^(1) = 2273.4, *p* < 0.001, v = 0.29, BF_10_ > 1000), meaning that even if they were not explicitly expecting another stimulus appearing besides the cross, the first detection question alone influenced detection rates. However, the detection rates in this trial were still reliably lower than in the control trial (Trial 8 - *χ*^2^(1) = 1449.4, *p* < 0.001, v = 0.23, BF_10_ > 1000).

To ensure that participants followed task instructions, we also analyzed performance on the cross task (see **Figure S2** – **Supplementary Material**). First, across trials, participants were able to perform the task above chance (M = 63.4%, SD = 18.3; t(13538) = 85.6, *p* < 0.001), suggesting that they were sufficiently engaged with it. There was also an overall difference between the trials, as shown by a one-way ANOVA with trial number as a within-subject factor (F(6.96, 94276.74) = 146.75, *p* < 0.001, η^2^p = 0.11). Notably, performance in the cross task was lower in Trial 7 (i.e., the divided attention trial, M = 64.32%, SD = 47.9) than in Trial 4, t(13538) = 4.84, *p* < 0.001, BF_10_ > 1000), likely because the presence of some expectation caused the stimulus appearing in the cross to be more distracting leading to a performance cost in the primary cross task. No such difference was observed when comparing Trial 4 (M = 67.61 %, SD = 46.8) and Trial 3 (M = 67.75%, SD = 46.7, t(13538) = 0.25, *p =* 0.800, BF_10_ = 0.01), confirming that participants continued to prioritize the cross task on Trial 4.

Together, these results show that we were able to reliably elicit inattentional blindness using the cross task on a large unsupervised sample of participants performing the task in a museum environment. As such, the detection rates we will be reporting for Trial 4 are informative of the bottom-up sensitivity towards the stimuli we have tested, under conditions of inattention. Detection rates for Trial 7, instead, likely reflect detection performance in the presence of some expectation towards a stimulus nearby the cross.

### 3.2 Trial 4 – No expectation

#### 3.2.1 Detection rates

In this trial, participants’ detection rates reflect detection performance without expectation of an additional stimulus beyond the cross. A logistic regression model on detection rates showed a main effect of Category (exp. β = 1.39, 95% CI [1.29, 1.45], *p* < 0.001, std. β = 8.87), with higher detection rates for persons than plants. The main effect of orientation was also significant (exp. β = 1.22, 95% CI [1.13, 1.31], *p* < 0.001, std. β = 5.33), with higher detection rates for upright than inverted stimuli. The logistic model also showed a main effect of Age (exp. β = 0.58, 95% CI [0.55, 0.60], *p* < 0.001, std. β = -28.94) and a main effect of reported Gender (exp. β = 0.76, 95% CI [0.71, 0.82], *p* < 0.001, std. β = -7.17), yet no models including the interaction between Age and Gender, nor models including the full interaction of these demographic factors and the Category x Orientation interaction significantly improved the explanatory power of the model (**See Supplementary Material** and section 3.4 for detailed analyses on the role of demographic variables on detection rates).

Crucially, the main effects of Orientation and Category were qualified by a significant Category x Orientation interaction (exp. β = 1.23, 95% CI [1.07, 1.43], *p* = 0.004, std. β = 2.85, **Figure 2A**). Using chi-squared tests we tested the inversion effect within each object category. The inversion effect was reliable for the detection rates of body stimuli (*χ*^2^(1) = 34.49, *p <* 0.001, v = 0.07, OR = 1.34, 95 % CI = [1.22, 1.48], BF_10_ > 1000). Upright bodies were detected by 61% of participants, while inverted bodies were detected by 53.7%. By contrast, the detection inversion effect was not reliably observed for plants (upright plant – 50.7%, inverted plant – 48.6 %; *χ*^2^(1) = 2.70, *p =* 0.100, v = 0.02, OR = 1.09, 95 % CI = [0.99, 1.20]), and the Bayesian analysis for this contrast indicated positive evidence for the null hypothesis (BF_10_ = 0.25).

**Figure 2.**
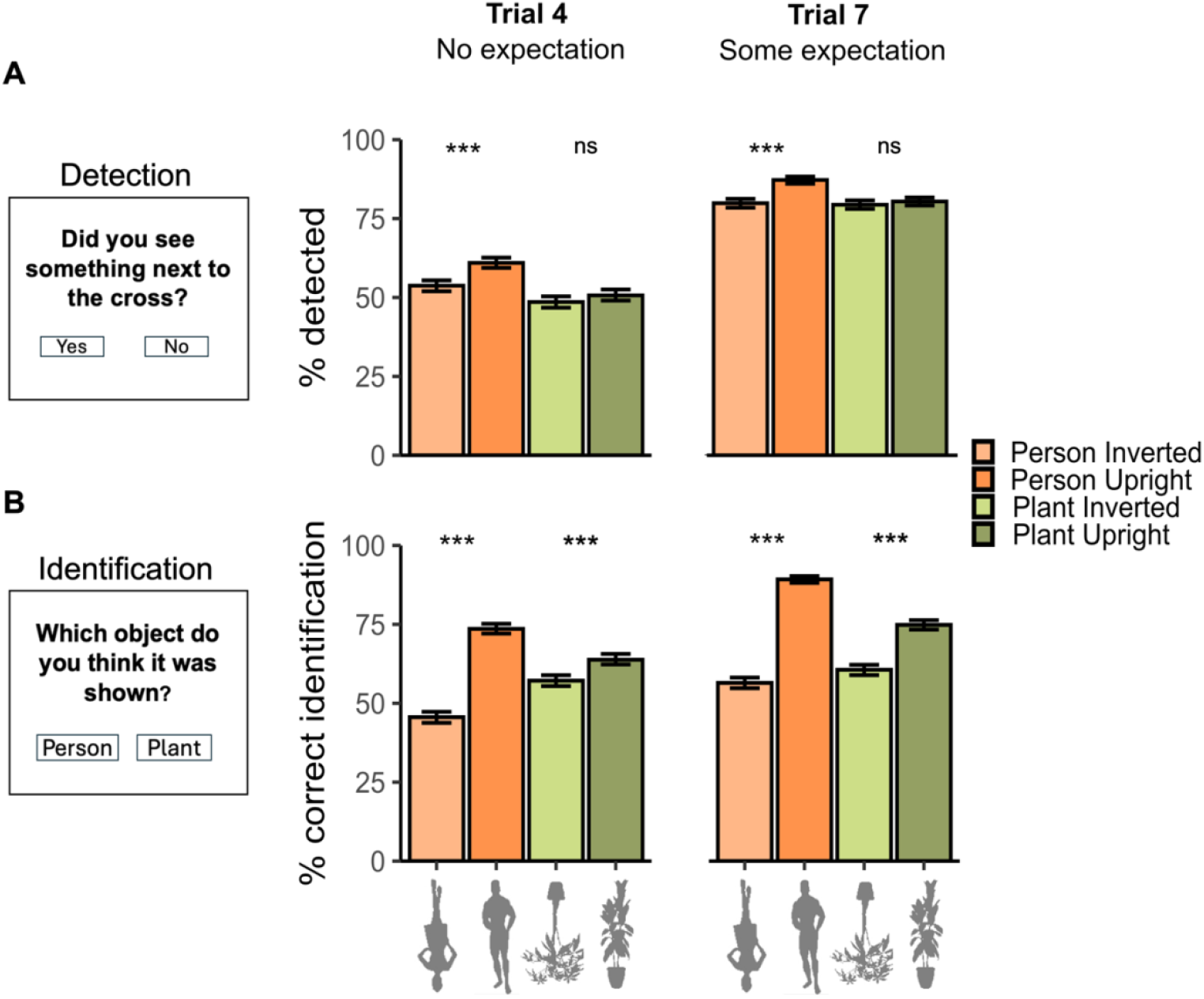
**A.** The bars show the observed average detection rates (in percentage) for each stimulus type in Trial 4 (on the left), when participants had no expectation of a stimulus appearing besides the cross, and in Trial 7 (on the right), when they had some expectation. **B**. The bars represent the percentage of correct identification for each object category and orientation in Trial 4 (on the left) and in Trial 7 (on the right). The error bars represent the 95% bootstrapped confidence interval of the mean. *** = p < 0.001; ns = Not significant. The paired comparisons refer to the chi-squared tests for upright vs inverted stimulus.

These results provide evidence for a detection inversion effect for person silhouettes. This effect could not be explained by top-down attention or expectation, and also not by differences in low-level features. Furthermore, the detection inversion effect was specific to bodies as it was absent for a control category (plants) and held even when regressing out overall detection differences due to age and reported gender.

#### 3.2.2 Identification performance

The identification question appeared right after the detection question. Participants were asked to identify the object, regardless of whether they had detected it (**Figure 1A**; see **Figure S5** for identification performance on the subset of trials where the stimulus had been detected, showing the same pattern of results). A logistic regression model on identification performance showed a main effect of Orientation (exp. β = 2.09, 95% CI [1.95, 2.25], *p* < 0.001, std. β = 20.01), and no main effect of Category (exp. β = 0.99, 95% CI [0.92, 1.07], *p* = 0.857, std. β = -0.18). Further, there was a main effect of Age (exp. β = 0.84, 95% CI [0.81, 0.88], *p* < 0.001, std. β = -9.18) and no significant main effect of reported Gender (exp. β = 0.97, 95% CI [0.90, 1.05], *p* < 0.001, std. β = -0.79). The interaction between Orientation and Category was also significant (exp. β = 2.52, 95% CI [2.18, 2.91], *p* < 0.001, std. β = 12.46, **Figure 2B**). We followed up on this interaction by testing the effect of inversion for each object category using chi-squared tests. Upright bodies were identified correctly more often (73.57% of the times) than their inverted counterpart (45.59%; *χ*^2^(1) = 530.88, *p <* 0.001, v = 0.30, OR = 3.32, 95% CI [2.99, 3.68], v = 0.29, BF_10_ > 1000). The inversion effect for the plant stimuli was smaller but significant (upright 63.79% vs inverted 57.16%, *χ*^2^(1) = 28.82, *p <* 0.001, OR = 1.35, 95 % CI [1.19, 1.46], v = 0.07, BF_10_ > 1000).

### 3.3 Trial 7 – Some Expectation

#### 3.3.1 Detection rates

At Trial 7, participants had previously been asked about having detected and identified another stimulus besides the cross and may thus have expected another stimulus to appear. Therefore, detection rates of the stimulus in this trial generally increase (see analysis across stimuli above). Yet, such expectations were not necessarily related to a specific object category or orientation. Indeed, by design, within the same participant the same object category and orientation combination was never repeated. Below and in **Figure 2A**, we report Trial 7 detection rates across all trials, while in **Figure S6** we report Trial 7 detection rates as a function of the stimulus that appeared in Trial 4 (results did not depend on the category shown in Trial 4).

A logistic regression model (see **Supplementary Material** for details on the selected model) on detection rates showed a main effect of Category (exp. β = 1.33, 95% CI [1.21, 1.46], *p* < 0.001, std. β = 5.93) and a main effect of Orientation (exp. β = 1.35, 95% CI [1.23, 1.48], *p* < 0.001, std. β = 6.26). The main effect of Age (exp. β = 0.57, 95% CI [0.55, 0.60], *p* < 0.001, std. β = -24.78), and Gender (exp. β = 0.81, 95% CI [0.74, 0.89], *p* < 0.001, std. β = -4.25) were also significant. Importantly, we also found a Category x Orientation interaction (exp. β = 1.67, 95% CI [1.38, 2.01], *p* < 0.001, std. β = 5.35, **Figure 2A**) indicative of a difference in inversion effects between each category. To further inspect this difference, we compared the detection rates in upright vs inverted conditions separately for people and plant stimuli using chi-squared tests. There was a significant inversion effect for person stimuli (*χ*^2^(1) = 62.30, *p <* 0.001, v = 0.10, OR = 1.72, 95 % CI = [1.50, 1.97], BF_10_ > 1000). Person stimuli were detected more often when presented upright (87.24%) than inverted (79.88%). The same inversion effect was not observed for the plant stimuli (upright - 79.85 %; inverted - 79.72%; χ^2^(1) = 1.01, *p =* 0.316, v = 0.01, OR = 1.07, 95% CI = [0.94, 1.20], BF_10_ = 0.08). These results provide further evidence for a person-specific detection inversion effect, even in the presence of *some* expectation for the presence of a stimulus.

#### 3.3.2 Identification performance

The logistic regression analysis on identification performance for Trial 7 revealed the same pattern as the analysis on identification performance for Trial 4. We again observed a main effect of Category (exp. β = 1.54, 95% CI [1.41, 1.68], *p* < 0.001, std. β = 9.93), a main effect of Orientation (exp. β = 3.54, 95% CI [3.25, 3.85], *p* < 0.001, std. β = 29.01), and a main effect of Age (exp. β = 0.80, 95% CI [0.77, 0.83], *p* < 0.001, std. β = -11.28), while there was no significant effect of Gender (exp. β = 1.00, 95% CI [0.92, 1.08], *p* = 0.97, std. β = -0.44). Furthermore, we again found a Category x Orientation interaction (exp. β = 3.36, 95% CI [2.83, 3.98], *p* < 0.001, std. β = 13.95, **Figure 2B**). Upright bodies were identified correctly more often (89.25%) than inverted bodies (56.46%, *χ*^2^(1) = 865.21, *p <* 0.001, v = 0.37, OR = 6.40, 95% CI = [5.61,7.31], BF_10_ > 1000). An inversion effect was also found for plants, although smaller in size (upright: 74.84%, inverted: 60.62%, *χ*^2^(1) = 149.13, *p <* 0.001, v = 0.15, OR = 1.93, 95% CI = [1.74, 2.15], BF_10_ > 1000).

### 3.4 The role of age and reported gender on inattentional blindness

#### 3.4.1 Detection rates

In the models reported above we regressed out the main effects of age and reported gender, showing that the Category x Orientation interaction was not modulated by these demographic variables. Effects of these demographic predictors have been previously reported, or incidentally observed, to modulate detection rates in inattentional blindness studies with relatively smaller sample sizes (age – Stothart et al., 2015; Graham and Burke, 2011; reported gender – Kreitz et al., 2015; Velichkovsky and Popova, 2020). While the primary aim of our study was not focused on revealing individual differences in inattentional blindness, we consider it useful to report and quantify such differences in our large sample, averaged across object category and orientation of the stimuli. In addition, our cross-task paradigm had two critical trials that differed in the expectancy of the participants for a stimulus (Trial 4 and Trial 7), such that we could assess if these individual differences are consistent across these trials.

We used logistic regression models including the following predictors: Age, Gender, and Trial (Trial 4 – no expectation/ Trial 7 – some expectation). The best model to explain detection rates (averaged across object category and orientation of the stimulus) included the three main effects of Age, Gender, and Trial. Including the interaction between Age and Gender, or between all three conditions, did not further improve the model fit (all *p*s from the likelihood ratio tests comparing the models with two- or three-way interactions to the main effect model were *p* > 0.283, see **Supplementary material**). For both trials (Trial 4 – M = 57.2%, W = 51.4%, Trial 7 – M = 83.3, W = 80.8), people identifying themselves as men were 1.22 times more likely to detect a stimulus than women (exp. β = 1.22, 95% CI [1.18, 1.27], *p* < 0.001, std. β = 8.36). Importantly, this gender difference was present despite the fact that performance of the two groups did not differ in the main cross task – (between subjects tests by gender in Trial 4 – *t*(12862) = 0.72, *p =* 0.473, BF_10_ = 0.03, Trial 7 – t(12862) = 0.59, *p* = 0.553, BF_10_ = 0.02, and on overall performance, t(12862) = -0.68, *p* = 0.498, BF_10_ = 0.03), suggesting that the increased inattentional blindness in women is not explained by gender differences in the primary cross task performance.

Concerning the age differences, the older were the participants the less likely they were to detect the stimulus appearing in the cross (exp. β = 0.60, 95% CI [0.58, 0.61], *p* < 0.001, std. β = -33.54). To facilitate visualization of the age differences, we plotted the model’s predicted probabilities of detecting the stimulus for both Trial 4 and Trial 7 (see **Figure 3A**). The probability of noticing a stimulus in the absence of expectation decreases on average by 7.45% (SD = 1.19) every 10 years of increased age. In the presence of some expectation (Trial 7), we observe an average decrease of 6.36% in detection every 10 years (SD = 1.97). Again, the age differences in the detection of the stimuli are not immediately explained by age differences in the main cross task, since we found no reliable correlation between age and the overall performance in the cross task (ρ = -01, *p* = 0.204, BF_10_ = 0.26).

**Figure 3.**
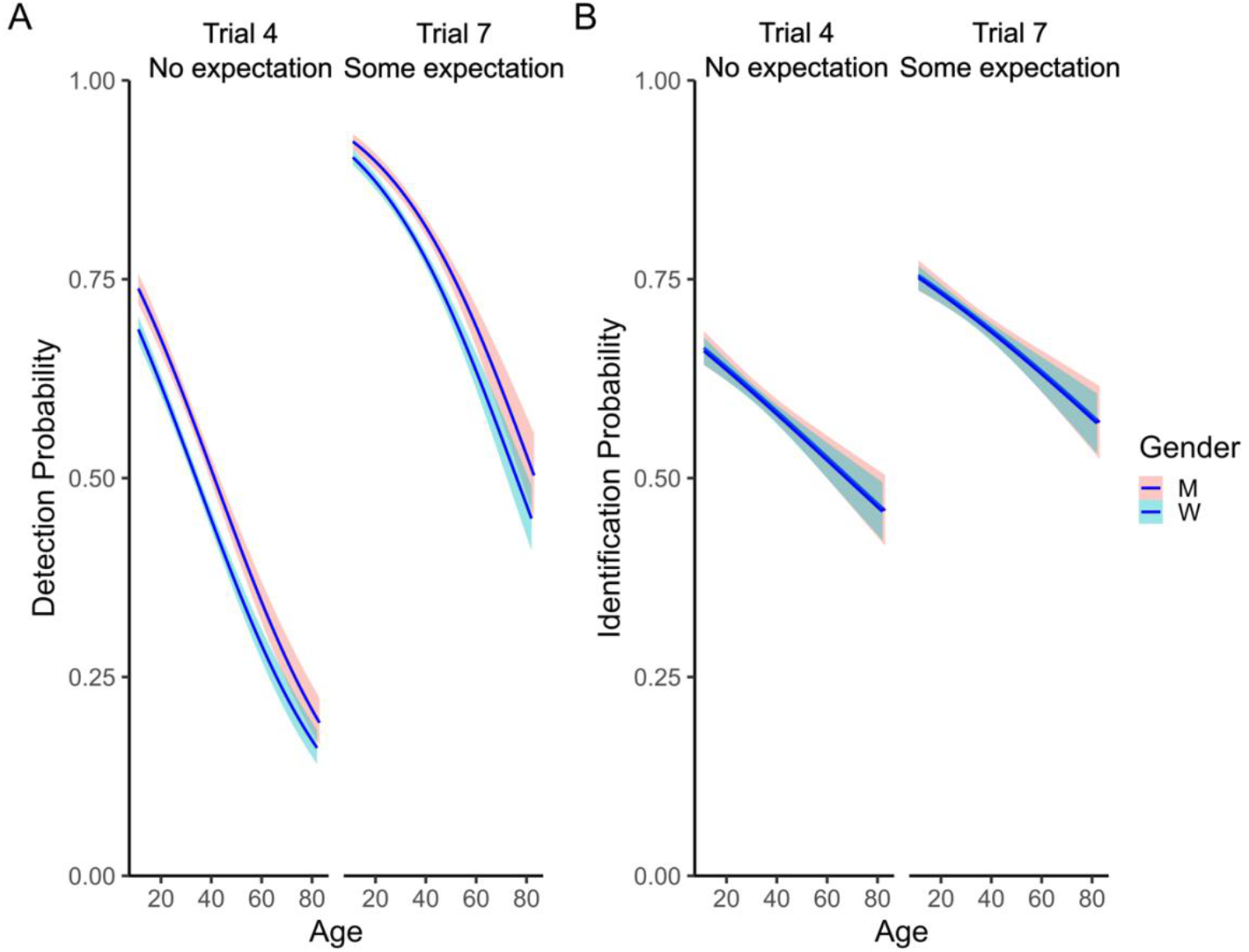
**A.** Estimated probability of detection based on the participants’ reported age and gender. The left panel shows the detection probability by age split between people identifying as women (light blue) and men (light red) in Trial 4 (No expectation). The right panel shows the estimate for Trial 7 (Some expectation). **B**. Estimated probability of identification based on the participants’ reported age and gender. The left panel shows the detection probability by age split between people identifying as women and men in Trial 4 (No expectation). The right panel shows the estimate for Trial 7 (Some expectation). The shaded area indicates the 95% confidence interval of the estimate.

#### 3.4.2 Identification performance

When testing the same model as above on identification performance (**Figure 3B**) we found a main effect of Age (exp. β = 0.83, 95% CI [0.81, 0.85], *p* < 0.001, std. β = -14.43). The probability of identifying the stimulus in absence of expectation (Trial 4) decreased on average by 2.57 % (SD = 0.21) every 10 years of increased age, while in Trial 7 by 2.85% (SD = 0.10). Differently from the detection rates, the participants’ gender did not reliably predict identification performance (exp. β = 0.98, 95% CI [0.93, 1.04], *p* = 0.528, std. β = - 0.63).

### 3.5 Headless bodies

The above results established a body detection inversion effect. It has been proposed that the body *recognition* inversion effect depends on the presence of the head. For example, studies using discrimination tasks showed that body inversion effects are generally weaker or even absent for headless bodies (Yovel et al., 2010; Brandman and Yovel, 2010; Axelsson et al., 2019; 2022; Robbins and Coltheart, 2012; Arizpe et al., 2017). To test whether the body detection inversion effect reported here similarly relies on the presence of the head, we ran an additional experiment using headless body silhouettes. For this experiment, we tested a new sample of N = 1327 participants (N = 328 participants with headless inverted bodies,N = 335 participants with headless upright bodies, N = 373 with inverted plants and N = 291 with upright plants) using the same set-up as used for the main experiment. Detailed methods and analyses are reported in the **Supplementary Material**. Briefly, we again found a difference between upright and inverted bodies, both in detection rates and identification performance, and both in Trial 4 and Trial 7 (**Figure 4 -** Trial 4 detection: *χ*^2^(1) = 7.11, *p* = 0.008, v = 0.11, OR = 1.54, 95 % CI = [1.13, 2.09], BF_10_ = 8.30; Trial 7 detection: *χ*^2^(1) = 6.44, *p* = 0.011, v = 0.11, OR = 1.81, 95 % CI = [1.16, 2.82], BF_10_ = 4.83; Trial 4 identification: *χ*^2^(1) = 41.28, *p* < 0.001, v = 0.25, OR = 2.83, 95 % CI = [2.06, 3.89], BF_10_ > 1000; Trial 7 identification: *χ*^2^(1) = 133.64, *p* < 0.001, v = 0.46, OR = 8.79, 95

**Figure 4.**
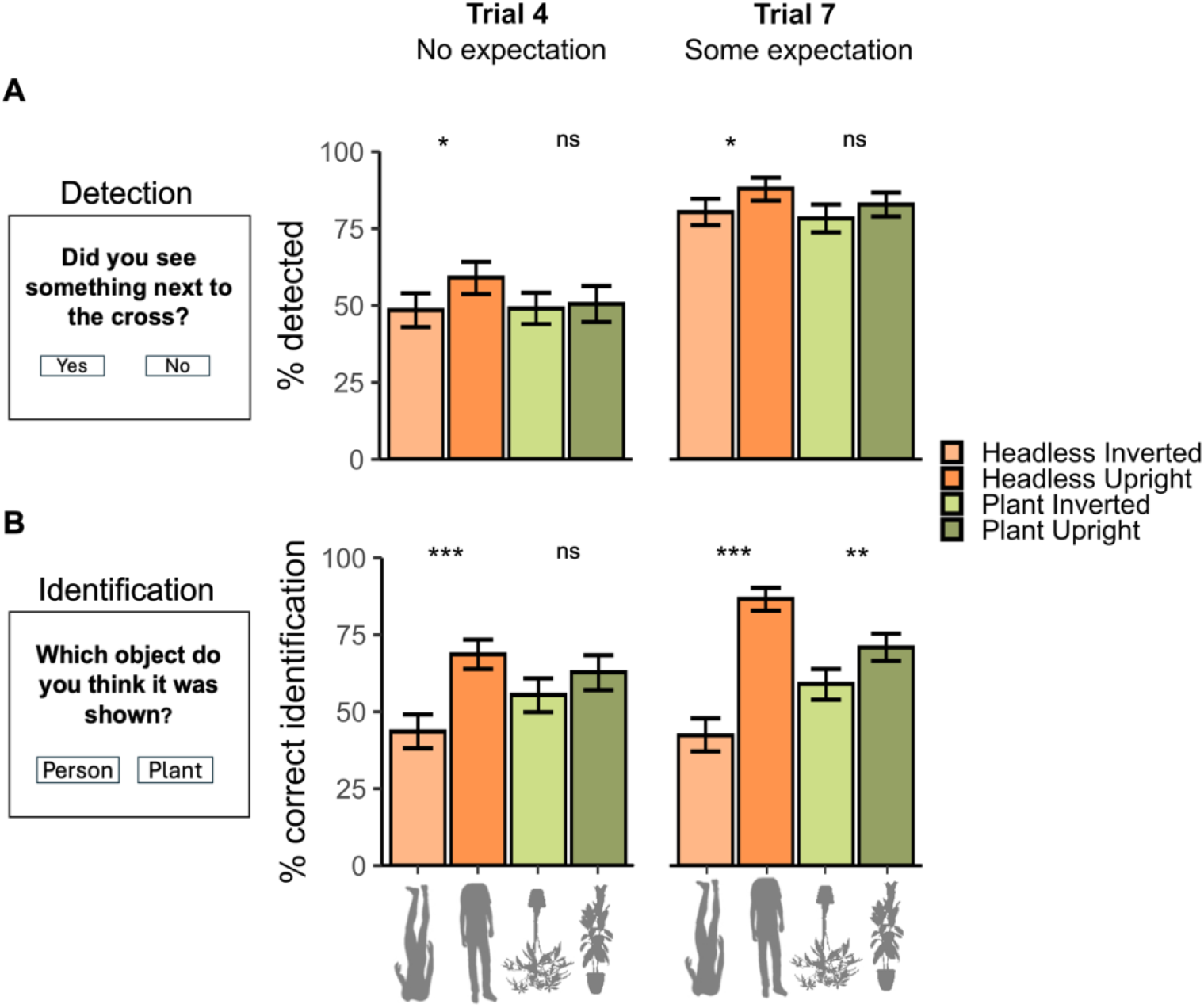
**A.** The bars show the observed average detection rates (in percentage) for upright and inverted headless bodies in Trial 4 (on the left), when participants had no expectation of a stimulus appearing besides the cross, and in Trial 7 (on the right), when they had some expectation. **B**. The bars represent the percentage of correct identification for upright and inverted headless bodies in Trial 4 (on the left) and in Trial 7 (on the right). The error bars represent the 95% bootstrapped confidence interval of the mean. * = p < 0.05; ** = p < 0.005; *** = p < 0.001. The paired comparisons refer to the chi-squared tests for upright vs inverted stimulus.

% CI = [5.92, 13.06], BF_10_ > 1000;). The detection inversion effect on Trial 4 for headless bodies in this experiment was comparable in size (10.6%) to the one found for full body silhouettes in the main experiment (7.3%; see **Supplementary Material** for analyses directly comparing detection and identification rates for bodies with and without heads), showing that the body detection inversion effect does not critically depend on the presence of the head.

## 4. Discussion

Through an inattentional blindness experiment in a large sample of naïve museum visitors, we found that human body stimuli were detected more often when upright than when inverted. The inversion effect during detection was specific for bodies and was not reliably observed for plant stimuli. This detection inversion effect emerged in conditions of “inattention”. That is, participants had no expectation of a body stimulus appearing on the screen and their attention was engaged in another, relatively challenging, visual task. The body-specific detection inversion effect replicated across critical Trial 4 and Trial 7. Trial 4 represents the purest measure of inattentional blindness, in which participants have no expectancy of *any* stimulus appearing on the screen besides the task-relevant cross. Instead, in Trial 7, participants have at least *some* expectation that another stimulus may appear in virtue of the detection question of Trial 4. However, due to the between-subject nature of the design, the same participant never saw the same stimulus (e.g., an upright person) in both Trial 4 and Trial 7, making it unlikely that the detection inversion effect observed in Trial 7 was driven by the expectancy of a body stimulus per se. As such, the results of Trial 4 and Trial 7 together provide converging and confirmatory evidence that detection systems are sensitive to the configuration of the human body.

A body-specific detection inversion effect under these conditions makes it unlikely that it reflects body-specific top-down attention (Thorat and Peelen, 2022). Rather, we interpret these findings as evidence that bottom-up detection mechanisms are sensitive to the configuration of the human body. An inversion effect emerging at the stage of stimulus detection suggests that the initial representation of the human body is configural, or at least not reducible to low-level features that distinguish body stimuli from other objects (e.g., luminance, spatial frequency, rectilinearity). However, our results leave open the question of what specific body features drive detection; specifically, whether it is driven by the whole-body structure (including torso, arms, and legs) or by smaller ensembles of body parts (e.g., arms linked to the torso, or the triangle formed by standing legs). For example, research coming from recognition tasks shows that behavioral effects indicative of configural processing are sensitive or to the symmetry between the limbs along the vertical midline of the body (R. A. Robbins & Coltheart, 2012a).

The body-specific inversion effect observed here and in previous work (using CFS; Stein et al., 2012) did not depend on the presence of the head, in contrast to what has been observed in body recognition tasks (e.g., identity or posture discrimination; Axelsson et al., 2019, 2022; Brandman & Yovel, 2010; R. A. Robbins & Coltheart, 2012c; Yovel et al., 2010). These previous studies had proposed that the head might be the main feature driving body inversion effects, and that these effects may therefore depend on face-specific rather than body-specific mechanisms. According to this view, body inversion effects would arise by virtue of strong contextual cueing of the body to a visible or implied face (Brandman & Yovel, 2010, 2012; Yovel et al., 2010). Different from body recognition, here we found that the body detection inversion effect was equally strong when bodies were presented with or without heads. This suggests that bottom-up detection mechanisms are tuned to the overall shape of the upright body (or subsets thereof), and that such tuning originates via body-specific rather than face-specific processing mechanisms.

What could be the neural mechanisms that underlie reduced inattentional blindness for upright bodies? Human and non-human primate research has shown that visual cortex contains regions that are selectively tuned to visual depictions of the body and its parts (Peelen & Downing, 2007; Vogels, 2022). There are several findings that suggest that this selectivity could drive the increased visual sensitivity to bodies under inattentional blindness, as observed here. First, body-selective regions are strongly and selectively activated by viewing minimal depictions of the body, including body silhouettes such as those used in the current study (Downing et al., 2001; Kana & Travers, 2012; Peelen & Downing, 2005; Schupp & Renner, 2011). Second, body-selective regions are selectively activated by body silhouettes that are task-irrelevant and presented outside the focus of attention (Thorat & Peelen, 2022), in line with bottom-up tuning to body shape. Third, brain stimulation studies have provided causal evidence for a role of the body-selective extrastriate body area (EBA; Downing et al., 2001) in the detection of bodies (van Koningsbruggen et al., 2013). Finally, while body-selective neurons in macaque visual cortex are tolerant to changes in viewpoint, position, and size, they are strongly sensitive to stimulus inversion (Popivanov et al., 2015). Altogether, these results suggest that the bottom-up visual sensitivity to upright depictions of the body, as observed here in human behavior, is mediated by neurons in visual cortex that are tuned to upright body shapes.

When participants were asked to recognize what object appeared on screen, *identification* performance for body stimuli was disproportionally affected by stimulus inversion, although inversion also impaired the recognition of plant stimuli. This finding is in line with previous results showing that inversion effects also occur for other familiar real-world objects, although smaller in size and less reliable than those observed with bodies or faces (e.g., Albonico et al., 2018; Gandolfo et al., 2024; Rezlescu et al., 2017; Robbins & McKone, 2007; Rossion & Curran, 2010). The reliable inversion effect for plant stimuli in identification but not in detection provides further evidence that identification and detection stages are dissociable, as previously shown by Mack et al. (2008). Recognizing an object as a plant likely depends on complex object features disrupted by inversion. By contrast, detecting a plant relies on simpler low-level visual features that are not disrupted by stimulus inversion.

The detection rates of the stimuli varied as a function of the object category and orientation of the stimuli inside the cross. Our design did not include a stimulus-absent condition in which no object is presented on Trial 4 but participants are still asked if they detected anything other than the cross. Previous studies implementing such a stimulus absent condition reported that participants sometimes falsely report the presence of a stimulus (Devue et al., 2009; Mack and Rock, 1998), perhaps because of a possible confusion of the stimulus with the following mask, or because of a decision strategy to “avoid appearing dumb or blind” (Mack and Rock 1998). It is therefore likely that in our study, across conditions, the detection rates were inflated. Future large-scale inattentional blindness studies may include a stimulus-absent condition to quantify the perceptual detection rates of the stimuli. Related to this, all the stimuli presented inside the cross were semantically meaningful (i.e., people and plants). Testing the detection rates of control stimuli such as visually matched mid-level textures (e.g., Grill-Spector and Kanwisher, 2005) could capture how meaningful familiar content generally affects detection performance in inattentional blindness (Devue et al., 2009; Mack and Rock, 1998).

Our findings contribute to the literature showing specialized processing of human bodies in various behavioral tasks. For example, during free viewing, participants preferentially look at the bodies and faces of people in natural scenes (Bindemann et al., 2010; Fletcher-Watson et al., 2008; Skripkauskaite et al., 2023). Body stimuli are also found more efficiently when they are the target of visual search, and they hinder search performance for other targets to a larger extent than other objects would (Langton et al., 2008; Ro et al., 2007). Extensive evidence shows that stimulus inversion disproportionally impairs fine-grained discrimination of those body characteristics that entail an understanding of the structural relations between parts such as posture, identity, sex (Arizpe et al., 2017; Axelsson et al., 2019, 2022; Gandolfo & Downing, 2020; Han et al., 2024; Minnebusch et al., 2010; Reed et al., 2003, 2006; R. A. Robbins & Coltheart, 2012b). That is, when presented in their typical, upright position, bodies are readily identified in virtue of the highly familiar relations among their single parts (e.g., arms on the sides of the torso, positioned above the legs). When inverted, these configural relations are disrupted, and recognition of these characteristics must be solved through part-based processing. Our results show that specialized visual processing of upright human bodies already occurs at the stage of detection, even in the absence of top-down influences.

### 4.1 Individual differences in Inattentional blindness

We acquired demographic information of age and gender of the participants. These demographic variables did not interact with the object category or orientation of the stimulus when predicting the detection rates (see **Supplementary Material**). However, they independently predicted the overall detection rates. People identifying as men detected the stimulus in Trial 4 and in Trial 7 more often than women did, and the probability of detecting the stimulus drastically decreased with an increase of age.

Previous research showed a similar negative relationship between age and detection rates. However, these studies demonstrated this with smaller samples, or by analyzing differences between different age groups without treating age as a continuous variable (Graham & Burke, 2011; Stothart et al., 2015). In addition, when sufficiently powered to detect individual differences, inattentional blindness tasks looking at aging effects were possibly influenced by sample self-selection (Stothart et al., 2015). Specifically, older adults approaching lab-testing or online testing platforms may have different attentional and cognitive abilities than the rest of the older population. The large-scale community-based sample of this study allowed to reduce, yet not fully eliminate, risks of sampling self-selection, and to bring a well-powered replication of these effects. It remains however an open question for future research to understand whether the age differences we reliably observed are specific to inattentional blindness (see also Stothart et al., 2015), or rather reflect a general decrease in attentional capacity occurring with ageing (Ball et al., 1988; Hasher & Zacks, 1988).

Gender effects - reduced susceptibility of men to inattentional blindness - were incidentally observed in a previous study analyzing the effect of personality variables on inattentional blindness (Kreitz, Schnuerch, et al., 2015) and studied systematically in one laboratory study with a much smaller sample (Velichkovsky & Popova, 2021). In this latter study, such gender differences did not depend on working memory load in the primary task. In line with this result, we observed no reliable gender differences in the performance of the primary cross task, suggesting that higher detection rates in men were not due to general differences in cognitive or perceptual abilities between men and women (Darda et al., 2020; Hyde, 2014; Miller & Halpern, 2014). Yet, the gender difference observed here must be carefully interpreted. For example, higher detection rates in men were not corroborated by higher accuracy in the identification of the stimuli in the identification question. Therefore, it is possible that the gender difference we observed in detection rates reflected more general differences in social behavior, in the decision criterion, or in the compliance with task instructions (Eagly, 1978, 1987; Eagly & Revelle, 2022), rather than differences in perceptual sensitivity. Future studies can disentangle these two possibilities by having a stimulus absent condition in each of the critical trials.

### 4.2 Testing single-trial attention paradigms in public spaces

The data of this study were collected as part of the Donders CityLab, a partnership between the Donders Institute and the muZIEum (https://muzieum.nl/), a museum about visual impairment and blindness. Collaborations between museums and researchers have previously been used for studies targeting specific populations (e.g., children; Callanan, 2012; Frank et al., 2012; Willard et al., 2019) or to assess the aesthetical experience of visitors with high ecological validity (Darda et al., 2024; Linden & Wagemans, 2024; Vissers & Wagemans, 2023). Here, we show that data collection on unsupervised participants in public spaces can generate robust results, improving our understanding of cognitive and perceptual phenomena. This approach is particularly useful for short (a few minutes) large-scale experiments which rely on single-trial measures in each participant. In our case, embedding the inattentional blindness paradigm in the context of an educational demonstration available to the public allowed us to reliably measure detection rates in the absence of expectation of any stimulus. In addition, collecting data in a museum gave us the chance to inform the public about perception and attention, thereby contributing to the museum’s mission. Specifically, we were able to raise awareness about inattentional blindness (i.e. cognitive blindness - Wolfe et al., 2022), a phenomenon relevant for daily-life tasks such as driving and sports (Charlton & Starkey, 2013; Furley et al., 2010; Xu et al., 2022).

### 4.3 Conclusions

The present study revealed that the human visual system is equipped with specialized mechanisms for detecting the presence of other people. A detection system tuned to upright depictions of the human body, irrespectively of the expectations and the attentional set of the observer, highlights the priority that the visual system assigns to detecting conspecifics. In natural viewing conditions, detection of another person is likely to occur from a distance. Bottom-up sensitivity to upright body features may provide an advantage, in evolutionary terms, for readily detecting the presence of others at a “safe” distance. Indeed, detecting others based on body cues may be the first step in the recognition of a potential threat as facial features may not be visible at a distance (Rice et al., 2013).

## Acknowledgements

We thank the members of the Donders Citylab project and the muZIEum staff for their support, and the Peelen Lab members for their feedback on this project.

## Open practices statement

The raw data generated from this project and the analysis code are publicly available at the Open Science Framework repository - https://osf.io/ahm7r/

## References

Albonico, A., Furubacke, A., Barton, J. J. S., & Oruc, I. (2018). Perceptual efficiency and the inversion effect for faces, words and houses. Vision Research, 153, 91–97. 10.1016/j.visres.2018.10.008

Aragon, T. J., Fay, M. P., Wollschlaeger, D., & Omidpanah, A. (2020). epitools: Epidemiology Tools (Version 0.5-10.1) [Computer software]. https://cran.r-project.org/web/packages/epitools/index.html

Arizpe, J. M., McKean, D. L., Tsao, J. W., & Chan, A. W.-Y. (2017). Where You Look Matters for Body Perception: Preferred Gaze Location Contributes to the Body Inversion Effect. PLOS ONE, 12(1), e0169148. 10.1371/journal.pone.0169148

Axelsson, E. L., Buddhadasa, T., Manca, L., & Robbins, R. A. (2022). Making heads or tails of body inversion effects: Do heads matter? PLOS ONE, 17(2), e0263902. 10.1371/journal.pone.0263902

Axelsson, E. L., Robbins, R. A., Copeland, H. F., & Covell, H. W. (2019). Body Inversion Effects With Photographic Images of Body Postures: Is It About Faces? Frontiers in Psychology, 10. https://www.frontiersin.org/articles/10.3389/fpsyg.2019.02686

Ball, K. K., Beard, B. L., Roenker, D. L., Miller, R. L., & Griggs, D. S. (1988). Age and visual search: Expanding the useful field of view. JOSA A, 5(12), 2210–2219. 10.1364/JOSAA.5.002210

Bao, H.-W.-S. (2024). bruceR: Broadly Useful Convenient and Efficient R Functions (Version 2024.6) [Computer software]. https://cran.r-project.org/web/packages/bruceR/index.html

Bindemann, M., Scheepers, C., Ferguson, H. J., & Burton, A. M. (2010). Face, body, and center of gravity mediate person detection in natural scenes. Journal of Experimental Psychology: Human Perception and Performance, 36(6), 1477–1485. 10.1037/a0019057

Brandman, T., & Yovel, G. (2010). The Body Inversion Effect Is Mediated by Face-Selective, Not Body-Selective, Mechanisms. Journal of Neuroscience, 30(31), 10534–10540. 10.1523/JNEUROSCI.0911-10.2010

Brandman, T., & Yovel, G. (2012). A face inversion effect without a face. Cognition, 125(3), 365–372. 10.1016/j.cognition.2012.08.001

Callanan, M. A. (2012). Conducting Cognitive Developmental Research in Museums: Theoretical Issues and Practical Considerations. Journal of Cognition and Development, 13(2), 137–151. 10.1080/15248372.2012.666730

Carey, S., De Schonen, S., Ellis, H. D., Bruce, V., Cowey, A., Ellis, A. W., & Perrett, D. I. (1992). Becoming a face expert. Philosophical Transactions of the Royal Society of London. Series B: Biological Sciences, 335(1273), 95–103. 10.1098/rstb.1992.0012

Cartwright-Finch, U., & Lavie, N. (2007). The role of perceptual load in inattentional blindness. Cognition, 102(3), 321–340. 10.1016/j.cognition.2006.01.002

Charlton, S. G., & Starkey, N. J. (2013). Driving on familiar roads: Automaticity and inattention blindness. Transportation Research Part F: Traffic Psychology and Behaviour, 19, 121–133. 10.1016/j.trf.2013.03.008

Darda, K. M., Butler, E. E., & Ramsey, R. (2020). Individual differences in social and non-social cognitive control. Cognition, 202, 104317. 10.1016/j.cognition.2020.104317

Darda, K. M., Gonzalez, V. E., Christensen, A. P., Bobrow, I., Krimm, A., Nasim, Z., Cardillo, E. R., Perthes, W., & Chatterjee, A. (2024). Engaging with art in-the-wild at the Barnes Foundation and Penn Museum. Research Square. 10.21203/rs.3.rs-4468529/v1

de la Rosa, S., Choudhery, R. N., & Chatziastros, A. (2011). Visual object detection, categorization, and identification tasks are associated with different time courses and sensitivities. Journal of Experimental Psychology: Human Perception and Performance, 37(1), 38–47. 10.1037/a0020553

de Leeuw, J. R. (2015). jsPsych: A JavaScript library for creating behavioral experiments in a Web browser. Behavior Research Methods, 47(1), 1–12. 10.3758/s13428-014-0458-y

Devue, C., Laloyaux, C., Feyers, D., Theeuwes, J., & Brédart, S. (2009). Do Pictures of Faces, and Which Ones, Capture Attention in the Inattentional-Blindness Paradigm? Perception, 38(4), 552–568. 10.1068/p6049

Downing, P. E., Bray, D., Rogers, J., & Childs, C. (2004). Bodies capture attention when nothing is expected. Cognition, 93(1), B27–B38. 10.1016/j.cognition.2003.10.010

Downing, P. E., Jiang, Y., Shuman, M., & Kanwisher, N. (2001). A cortical area selective for visual processing of the human body. Science (New York, N.Y.), 293(5539), 2470–2473. 10.1126/science.1063414

Eagly, A. H. (1978). Sex differences in influenceability. Psychological Bulletin, 85(1), 86–116. 10.1037/0033-2909.85.1.86

Eagly, A. H. (1987). Sex Differences in Social Behavior: A Social-role interpretation. Psychology Press. 10.4324/9780203781906

Eagly, A. H., & Revelle, W. (2022). Understanding the Magnitude of Psychological Differences Between Women and Men Requires Seeing the Forest and the Trees. Perspectives on Psychological Science, 17(5), 1339–1358. 10.1177/17456916211046006

Farah, M. J., Tanaka, J. W., & Drain, H. M. (1995). What causes the face inversion effect? Journal of Experimental Psychology: Human Perception and Performance, 21(3), 628–634. 10.1037/0096-1523.21.3.628

Farah, M. J., Wilson, K. D., Drain, M., & Tanaka, J. N. (1998). What is “special” about face perception? Psychological Review, 105(3), 482–498. 10.1037/0033-295X.105.3.482

Fletcher-Watson, S., Findlay, J. M., Leekam, S. R., & Benson, V. (2008). Rapid Detection of Person Information in a Naturalistic Scene. Perception, 37(4), 571–583. 10.1068/p5705

Frank, M. C., Vul, E., & Saxe, R. (2012). Measuring the Development of Social Attention Using Free-Viewing. Infancy, 17(4), 355–375. 10.1111/j.1532-7078.2011.00086.x

Furley, P., Memmert, D., & Heller, C. (2010). The dark side of visual awareness in sport: Inattentional blindness in a real-world basketball task. Attention, Perception, & Psychophysics, 72(5), 1327–1337. 10.3758/APP.72.5.1327

Gandolfo, M., Abassi, E., Balgova, E., Downing, P. E., Papeo, L., & Koldewyn, K. (2024). Converging evidence that left extrastriate body area supports visual sensitivity to social interactions. Current Biology, 34(2), 343-351.e5. 10.1016/j.cub.2023.12.009

Gandolfo, M., & Downing, P. E. (2020). Asymmetric visual representation of sex from human body shape. Cognition, 205, 104436. 10.1016/j.cognition.2020.104436

Graham, E. R., & Burke, D. M. (2011). Aging increases inattentional blindness to the gorilla in our midst. Psychology and Aging, 26(1), 162–166. 10.1037/a0020647

Grill-Spector, K., & Kanwisher, N. (2005). Visual Recognition: As Soon as You Know It Is There, You Know What It Is. Psychological Science, 16(2), 152–160. 10.1111/j.0956-7976.2005.00796.x

Han, Q., Gandolfo, M., & Peelen, M. V. (2024). Prior knowledge biases the visual memory of body postures. iScience, 27(4). 10.1016/j.isci.2024.109475

Hannon, E. M., & Richards, A. (2010). Is Inattentional Blindness Related to Individual Differences in Visual Working Memory Capacity or Executive Control Functioning? Perception, 39(3), 309–319. 10.1068/p6379

Hasher, L., & Zacks, R. T. (1988). Working Memory, Comprehension, and Aging: A Review and a New View. In G. H. Bower (Ed.), Psychology of Learning and Motivation (Vol. 22, pp. 193–225). Academic Press. 10.1016/S0079-7421(08)60041-9

Hershler, O., & Hochstein, S. (2005). At first sight: A high-level pop out effect for faces. Vision Research, 45(13), 1707–1724. 10.1016/j.visres.2004.12.021

Hutchinson, B. T., Pammer, K., Bandara, K., & Jack, B. N. (2022). A tale of two theories: A meta-analysis of the attention set and load theories of inattentional blindness. Psychological Bulletin, 148(5–6), 370–396. 10.1037/bul0000371

Hyde, J. S. (2014). Gender Similarities and Differences. Annual Review of Psychology, 65(Volume 65, 2014), 373–398. 10.1146/annurev-psych-010213-115057

Jeffreys, H. (1961). The Theory of Probability. OUP Oxford.

Jiang, Y., Costello, P., & He, S. (2007). Processing of Invisible Stimuli: Advantage of Upright Faces and Recognizable Words in Overcoming Interocular Suppression. Psychological Science, 18(4), 349–355. 10.1111/j.1467-9280.2007.01902.x

Kana, R. K., & Travers, B. G. (2012). Neural substrates of interpreting actions and emotions from body postures. Social Cognitive and Affective Neuroscience, 7(4), 446–456. 10.1093/scan/nsr022

Koningsbruggen, M. G. van, Peelen, M. V., & Downing, P. E. (2013). A Causal Role for the Extrastriate Body Area in Detecting People in Real-World Scenes. Journal of Neuroscience, 33(16), 7003–7010. 10.1523/JNEUROSCI.2853-12.2013

Kreitz, C., Furley, P., Memmert, D., & Simons, D. J. (2015). Inattentional Blindness and Individual Differences in Cognitive Abilities. PLOS ONE, 10(8), e0134675. 10.1371/journal.pone.0134675

Kreitz, C., Schnuerch, R., Gibbons, H., & Memmert, D. (2015). Some See It, Some Don’t: Exploring the Relation between Inattentional Blindness and Personality Factors. PLOS ONE, 10(5), e0128158. 10.1371/journal.pone.0128158

Kuroki, D. (2021). A new jsPsych plugin for psychophysics, providing accurate display duration and stimulus onset asynchrony. Behavior Research Methods, 53(1), 301–310. 10.3758/s13428-020-01445-w

Langton, S. R. H., Law, A. S., Burton, A. M., & Schweinberger, S. R. (2008). Attention capture by faces. Cognition, 107(1), 330–342. 10.1016/j.cognition.2007.07.012

Lee, M. D., & Wagenmakers, E.-J. (2014). Bayesian Cognitive Modeling: A Practical Course. Cambridge University Press.

Linden, C., & Wagemans, J. (2024). Presenting TaMuNaBe: A taxonomy of museum navigation behaviors. Psychology of Aesthetics, Creativity, and the Arts, 18(2), 143–159. 10.1037/aca0000413

Long, J. A. (2024). jtools: Analysis and Presentation of Social Scientific Data. Journal of Open Source Software, 9(101), 6610. 10.21105/joss.06610

Ly, A., Verhagen, J., & Wagenmakers, E.-J. (2016). Harold Jeffreys’s default Bayes factor hypothesis tests: Explanation, extension, and application in psychology. Journal of Mathematical Psychology, 72, 19–32. 10.1016/j.jmp.2015.06.004

Mack, A., & Rock, I. (1998). Inattentional blindness (pp. xiv, 273). The MIT Press.

Mack, M. L., Gauthier, I., Sadr, J., & Palmeri, T. J. (2008). Object detection and basic-level categorization: Sometimes you know it is there before you know what it is. Psychonomic Bulletin & Review, 15(1), 28–35. 10.3758/PBR.15.1.28

Miller, D. I., & Halpern, D. F. (2014). The new science of cognitive sex differences. Trends in Cognitive Sciences, 18(1), 37–45. 10.1016/j.tics.2013.10.011

Minnebusch, D. A., Keune, P. M., Suchan, B., & Daum, I. (2010). Gradual inversion affects the processing of human body shapes. NeuroImage, 49(3), 2746–2755. 10.1016/j.neuroimage.2009.10.046

Morey, R. D., Rouder, J. N., Jamil, T., Urbanek, S., Forner, K., & Ly, A. (2024). BayesFactor: Computation of Bayes Factors for Common Designs (Version 0.9.12-4.7) [Computer software]. https://cran.r-project.org/web/packages/BayesFactor/index.html

Nakayama, K., He, Z. J., & Shimojo, S. (1995). Visual Surface Representation: A Critical Link between Lower-Level and Higher-Level Vision. 10.7551/mitpress/3965.003.0004

Paffen, C. L. E., & Alais, D. (2011). Attentional modulation of binocular rivalry. Frontiers in Human Neuroscience, 5. 10.3389/fnhum.2011.00105

Pedersen, T. L. (2024). patchwork: The Composer of Plots (Version 1.3.0) [Computer software]. https://cran.r-project.org/web/packages/patchwork/index.html

Peelen, M. V., & Downing, P. E. (2005). Selectivity for the Human Body in the Fusiform Gyrus. Journal of Neurophysiology, 93(1), 603–608. 10.1152/jn.00513.2004

Peelen, M. V., & Downing, P. E. (2007). The neural basis of visual body perception. Nature Reviews. Neuroscience, 8(8), 636–648. 10.1038/nrn2195

Popivanov, I. D., Jastorff, J., Vanduffel, W., & Vogels, R. (2015). Tolerance of Macaque Middle STS Body Patch Neurons to Shape-preserving Stimulus Transformations. Journal of Cognitive Neuroscience, 27(5), 1001–1016. 10.1162/jocn_a_00762

R: The R Project for Statistical Computing. (n.d.). Retrieved October 4, 2024, from https://www.r-project.org/

Reed, C. L., Stone, V. E., Bozova, S., & Tanaka, J. (2003). The Body-Inversion Effect. Psychological Science, 14(4), 302–308. 10.1111/1467-9280.14431

Reed, C. L., Stone, V. E., Grubb, J. D., & McGoldrick, J. E. (2006). Turning configural processing upside down: Part and whole body postures. Journal of Experimental Psychology: Human Perception and Performance, 32(1), 73–87. 10.1037/0096-1523.32.1.73

Remington, A., Cartwright-Finch, U., & Lavie, N. (2014). I can see clearly now: The effects of age and perceptual load on inattentional blindness. Frontiers in Human Neuroscience, 8. 10.3389/fnhum.2014.00229

Rezlescu, C., Susilo, T., Wilmer, J. B., & Caramazza, A. (2017). The inversion, part-whole, and composite effects reflect distinct perceptual mechanisms with varied relationships to face recognition. Journal of Experimental Psychology: Human Perception and Performance, 43(12), 1961–1973. 10.1037/xhp0000400

Rice, A., Phillips, P. J., Natu, V., An, X., & O’Toole, A. J. (2013). Unaware Person Recognition From the Body When Face Identification Fails. Psychological Science, 24(11), 2235–2243. 10.1177/0956797613492986

Richards, A., Hannon, E. M., & Derakshan, N. (2010). Predicting and manipulating the incidence of inattentional blindness. Psychological Research, 74(6), 513–523. 10.1007/s00426-009-0273-8

Ro, T., Friggel, A., & Lavie, N. (2007). Attentional biases for faces and body parts. Visual Cognition, 15(3), 322–348. 10.1080/13506280600590434

Robbins, R. A., & Coltheart, M. (2012a). Left-right holistic integration of human bodies. Quarterly Journal of Experimental Psychology (2006), 65(10), 1962–1974. 10.1080/17470218.2012.674145

Robbins, R. A., & Coltheart, M. (2012b). The effects of inversion and familiarity on face versus body cues to person recognition. Journal of Experimental Psychology: Human Perception and Performance, 38(5), 1098–1104. 10.1037/a0028584

Robbins, R. A., & Coltheart, M. (2012c). The effects of inversion and familiarity on face versus body cues to person recognition. Journal of Experimental Psychology: Human Perception and Performance, 38(5), 1098–1104. 10.1037/a0028584

Robbins, R., & McKone, E. (2007). No face-like processing for objects-of-expertise in three behavioural tasks. Cognition, 103(1), 34–79. 10.1016/j.cognition.2006.02.008

Rosch, E., Mervis, C. B., Gray, W. D., Johnson, D. M., & Boyes-Braem, P. (1976). Basic objects in natural categories. Cognitive Psychology, 8(3), 382–439. 10.1016/0010-0285(76)90013-X

Rossion, B., & Curran, T. (2010). Visual Expertise with Pictures of Cars Correlates with RT Magnitude of the Car Inversion Effect. Perception, 39(2), 173–183. 10.1068/p6270

Schupp, H. T., & Renner, B. (2011). The Implicit Nature of the Anti-Fat Bias. Frontiers in Human Neuroscience, 5. 10.3389/fnhum.2011.00023

Skripkauskaite, S., Mihai, I., & Koldewyn, K. (2023). Attentional bias towards social interactions during viewing of naturalistic scenes. Quarterly Journal of Experimental Psychology, 76(10), 2303–2311. 10.1177/17470218221140879

Stein, T. (2019). The Breaking Continuous Flash Suppression Paradigm: Review, evaluation, and outlook. In Transitions Between Consciousness and Unconsciousness. Routledge.

Stein, T., & Peelen, M. V. (2015). Content-specific expectations enhance stimulus detectability by increasing perceptual sensitivity. Journal of Experimental Psychology: General, 144(6), 1089–1104. 10.1037/xge0000109

Stein, T., & Peelen, M. V. (2021). Dissociating conscious and unconscious influences on visual detection effects. Nature Human Behaviour, 5(5), 612–624. 10.1038/s41562-020-01004-5

Stein, T., Sterzer, P., & Peelen, M. V. (2012). Privileged detection of conspecifics: Evidence from inversion effects during continuous flash suppression. Cognition, 125(1), 64–79. 10.1016/j.cognition.2012.06.005

Stewart, N., Chandler, J., & Paolacci, G. (2017). Crowdsourcing Samples in Cognitive Science. Trends in Cognitive Sciences, 21(10), 736–748. 10.1016/j.tics.2017.06.007

Störmer, V. S., Cohen, M. A., & Alvarez, G. A. (2019). Tuning Attention to Object Categories: Spatially Global Effects of Attention to Faces in Visual Processing. Journal of Cognitive Neuroscience, 31(7), 937–947. 10.1162/jocn_a_01400

Stothart, C. R., Boot, W. R., & Simons, D. J. (2015). Using Mechanical Turk to Assess the Effects of Age and Spatial Proximity on Inattentional Blindness. Collabra, 1(1), 2. 10.1525/collabra.26

Thorat, S., & Peelen, M. V. (2022). Body shape as a visual feature: Evidence from spatially-global attentional modulation in human visual cortex. NeuroImage, 255, 119207. 10.1016/j.neuroimage.2022.119207

Tsuchiya, N., & Koch, C. (2005). Continuous flash suppression reduces negative afterimages. Nature Neuroscience, 8(8), 1096–1101. 10.1038/nn1500

Velichkovsky, B. B., & Popova, S. (2021). Gender Differences in Object and Spatial Inattentional Blindness Under Working Memory Load. In B. M. Velichkovsky, P. M. Balaban, & V. L. Ushakov (Eds.), Advances in Cognitive Research, Artificial Intelligence and Neuroinformatics (pp. 122–127). Springer International Publishing. 10.1007/978-3-030-71637-0_14

Vissers, N., & Wagemans, J. (2023). Beyond the single picture: Aesthetic experiences with photography series in an exhibition context. Psychology of Aesthetics, Creativity, and the Arts, 17(5), 619–631. 10.1037/aca0000417

Vogels, R. (2022). More Than the Face: Representations of Bodies in the Inferior Temporal Cortex. Annual Review of Vision Science, 8(1), 383–405. 10.1146/annurev-vision-100720-113429

Wickham, H., Averick, M., Bryan, J., Chang, W., McGowan, L. D., François, R., Grolemund, G., Hayes, A., Henry, L., Hester, J., Kuhn, M., Pedersen, T. L., Miller, E., Bache, S. M., Müller, K., Ooms, J., Robinson, D., Seidel, D. P., Spinu, V., … Yutani, H. (2019). Welcome to the Tidyverse. Journal of Open Source Software, 4(43), 1686. 10.21105/joss.01686

Willard, A. K., Busch, J. T. A., Cullum, K. A., Letourneau, S. M., Sobel, D. M., Callanan, M., & Legare, C. H. (2019). Explain This, Explore That: A Study of Parent–Child Interaction in a Children’s Museum. Child Development, 90(5), e598–e617. 10.1111/cdev.13232

Wolfe, J. M., Kosovicheva, A., & Wolfe, B. (2022). Normal blindness: When we Look But Fail To See. Trends in Cognitive Sciences, 26(9), 809–819. 10.1016/j.tics.2022.06.006

Xu, J., Park, S. H., Zhang, X., & Hu, J. (2022). The Improvement of Road Driving Safety Guided by Visual Inattentional Blindness. IEEE Transactions on Intelligent Transportation Systems, 23(6), 4972–4981. IEEE Transactions on Intelligent Transportation Systems. 10.1109/TITS.2020.3044927

Yin, R. K. (1969). Looking at upside-down faces. Journal of Experimental Psychology, 81(1), 141–145. 10.1037/h0027474

Yovel, G., Pelc, T., & Lubetzky, I. (2010). It’s all in your head: Why is the body inversion effect abolished for headless bodies? Journal of Experimental Psychology: Human Perception and Performance, 36(3), 759–767. 10.1037/a0017451

