## Supplementary Material for "A body detection inversion effect revealed by a large-scale inattentional blindness experiment"

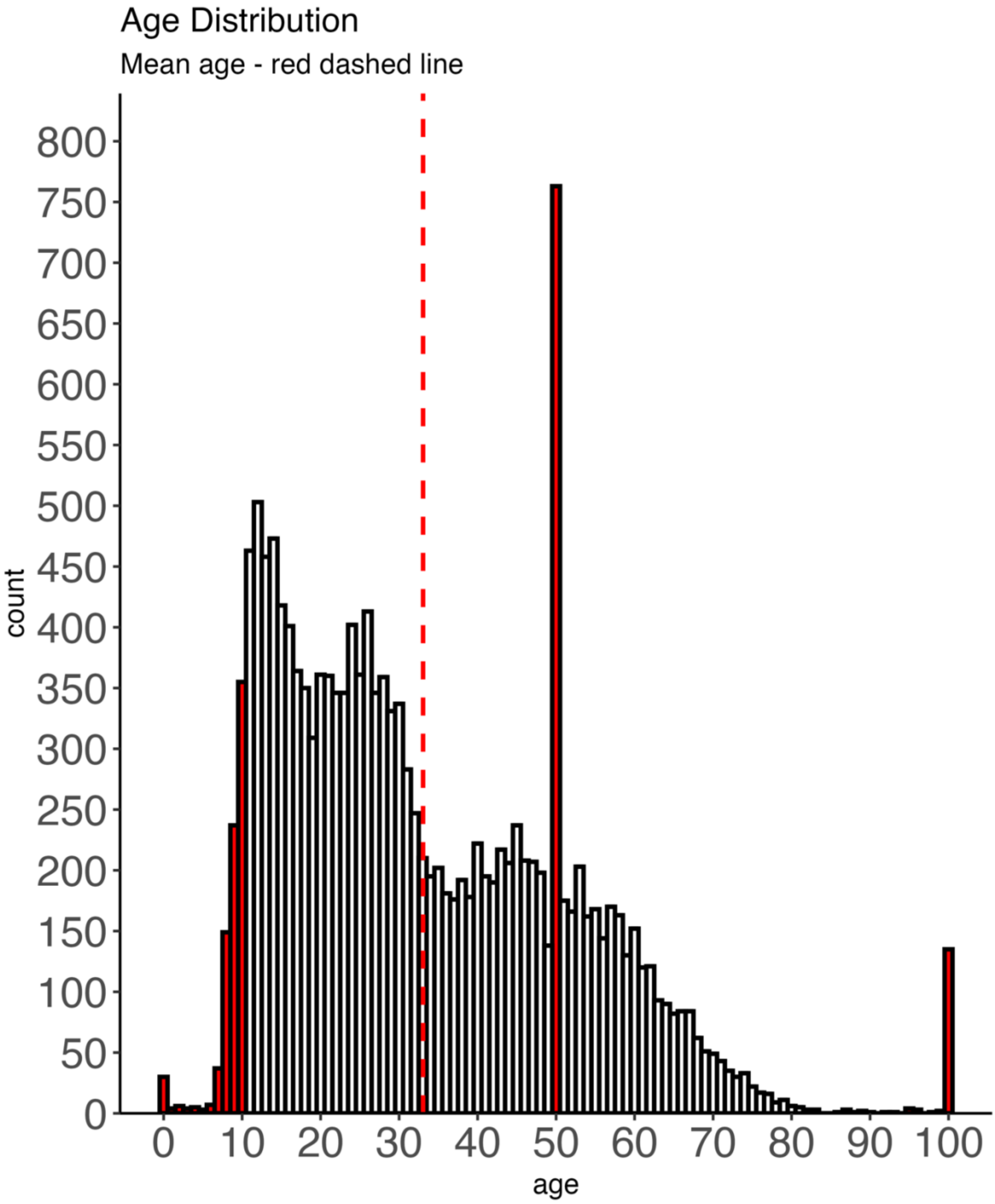


**Figure S1**. *Histogram for the age distribution in the population sample. The red dashed line represents the mean age of the whole distribution. The red histograms' bars represent the excluded datafiles based on age. For example, the spike of 50 years old reflects the default choice on the slider, leading to unreliable age reporting for the 50 years' old participants.*

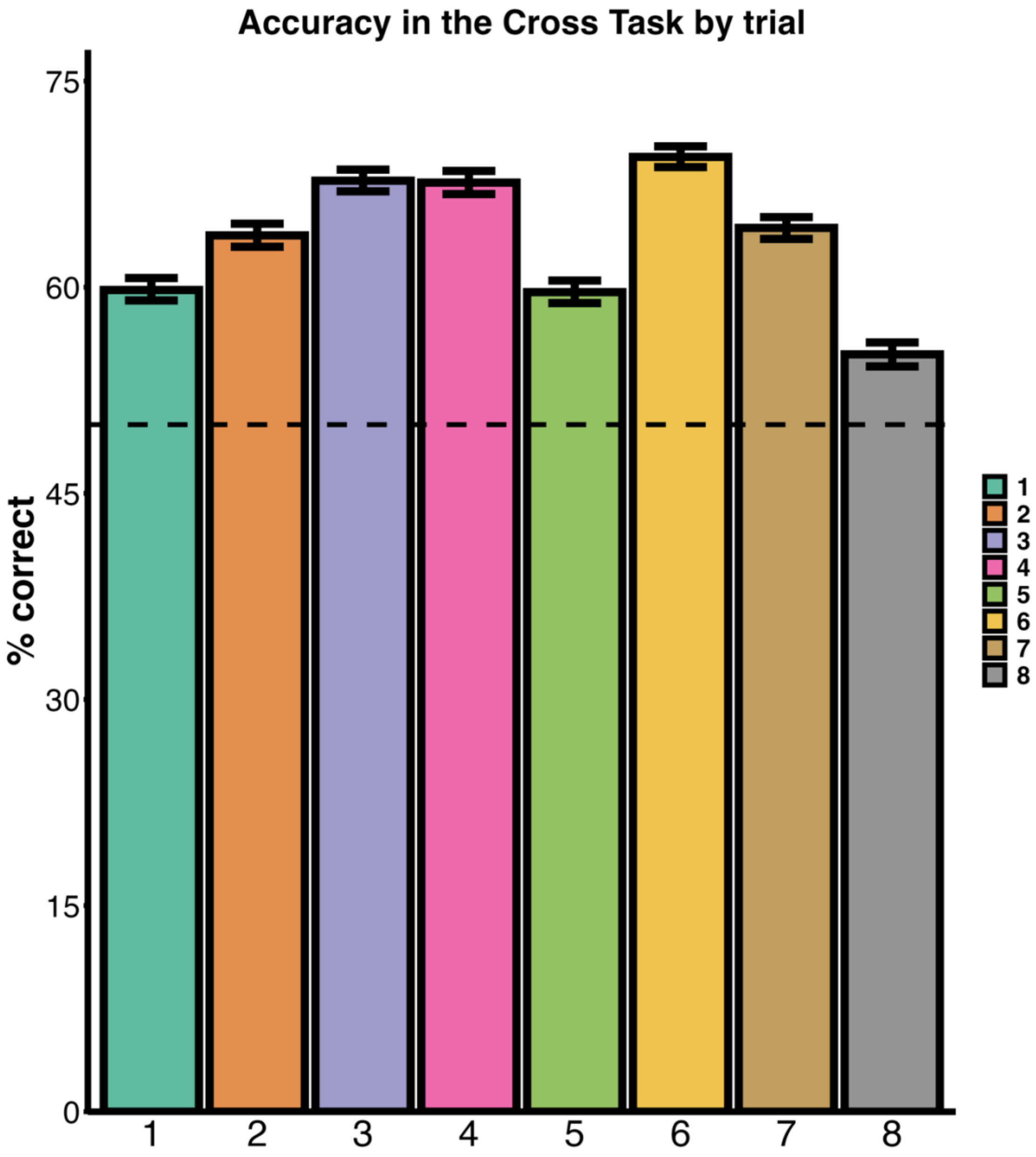


**Figure S2.** *Accuracy in the cross-task in each trial from 1-8. Participant's accuracy was overall above chance. On trial 4, when the stimulus appeared, performance was not affected as compared to trial 3, meaning that participants were performing the critical trial 4 without expectations about upcoming stimuli. Performance instead dropped in trial 5, right after they were presented with the first detection question, suggesting that participants divided their attention following the question related to the stimulus and looked for a stimulus besides the cross. In trial 7, performance of the cross task because, attention was diverted to the already expected stimulus, indeed, performance dropped as compared to Trial 4. In Trial 8,*

*performance furtherly dropped because participants were asked explicitly to focus on identifying the stimulus appearing besides the cross, rather than on the cross task itself.*

**Model selection procedure for the stimulus x orientation analyses**

We adopted a stepwise logistic regression approach with forward selection (Stoltzfus, 2011; Darlington, 1990; Hosmer et al., 2013) of the predictors and ran models predicting either detection rates, or identification scores separately for Trial 4 and Trial 7. The predictors were the Object Category (Person/Plant), the Orientation (Upright/Inverted), the Age, and the Gender of the participants. The Object Category and the Orientation predictors were contrast coded (the grand mean of the levels of the contrast coded predictor is taken as the intercept), while the Gender was dummy coded. Model performance for selection was assessed by performing a likelihood ratio test between the simpler and more complex model, and by looking at the Akaike Information Criterion (AIC – Bozdogan, 1987; Akaike, 1981), a measure that penalizes the addition of factors. If an additional predictor, or the interaction term among predictors, improved model fit significantly this was included in the model. We started by including the main effect of Category and of Orientation, then tested whether adding the interaction term between these predictors improved model fit. Further, we assessed whether including the main effect of age and gender, or the interaction, or an overall interaction between all the factors improved the model performance (see Table S1).

Table S1.

*Models' Performance for detection rates in Trial 4.*

| ***Model*** | ***AIC*** | ***P.Value(Chi$^2$)*** |
|---|---|---|
| **Det ~ Object Category + Orientation** | 17664.5 | |
| **Det ~ Object Category*Orientation** | 17657.6 | 0.003 |
| **Det ~ Object Category*Orientation + Age + Gender** | **16722.6** | **< 0.001** |
| **Det ~ Object Category*Orientation + Age*Gender** | 16724.3 | 0.62 |
| **Det ~ Object Category*Orientation*Age*Gender** | 16736.6 | 0.77 |

*Models' Performance for detection rates in Trial 7.*

| ***Model*** | ***AIC*** | ***P.Value(Chi$^2$)*** |
|---|---|---|

| | | |
|---|---|---|
| **Det ~ Object Category + Orientation** | 12180.7 | |
| **Det ~ Object Category*Orientation** | 12155.9 | < 0.001 |
| **Det ~ Object Category*Orientation + Age + Gender** | **11520.1** | **< 0.001** |
| **Det ~ Object Category*Orientation + Age*Gender** | 11521.3 | 0.359 |
| **Det ~ Object Category*Orientation*Age*Gender** | 11531.5 | 0.567 |

*Note.* The selected model reported in the main results' section is highlighted in bold font. The p-values compare the more complex models to the simpler model which significantly improved model fit.

**Model selection procedure for the demographic variables analysis**

The demographic variables significantly predicted the detection rates in Trial 4 and Trial 7 independently of the Category and Orientation. We ran separate models considering as predictors the age and gender variables on detection rates (see Table S2). In addition, for these models we also included as predictor the Trial – (Trial 4 and Trial 7), to exploratorily assess whether the individual differences of age and gender were directly modulated by the presence of some vs no-expectations for detecting a stimulus. The model explaining best the detection rates was the model including the three main effects of Age, Gender, and Trial. The probability of detecting the stimulus decreased with age, and was increased for man. The amount of expectations for a stimulus increased the detection rates irrespectively of the age and gender of the participants.

Table S2.
*Models' Performance testing demographic variables on detection rates.*

| ***Model*** | ***AIC*** | ***P.Value(Chi$^2$)*** |
|---|---|---|
| **Det ~ Age** | 31031.4 | |
| **Det ~ Age + Gender** | 30969.6 | < 0.001 |
| **Det ~ Age + Gender + Trial** | **28437.08** | **< 0.001** |
| **Det ~ Age*Gender + Trial** | 28437.9 | 0.28 |

| | | |
|---|---|---|
| **Det ~ Age*Trial + Gender** | 28439.0 | 0.79 |
| **Det ~ Gender*Trial + Age** | 28438.1 | 0.32 |
| **Det ~ Age*Gender*Trial** | 28443.1 | 0.75 |

*Note.* The selected model reported in the main results' section is highlighted in bold font. The p-values compare the more complex models to simpler best fitting models.

**Analyses on Trial 8**

Results

*Detection rates*

The results of the detection rates are shown in Figure S3A. Here, we tested the same logistic regression models as in Trial 4 and Trial 7 presented in the main text. The model on detection rates showed a main effect of Orientation (exp. $\beta$ = 1.50, 95% CI [1.27, 1.77], $p <$ 0.001, std. $\beta$ = 4.85), with higher detection rates for upright than inverted stimuli and a main effect of Object Category (exp. $\beta$ = 1.47, 95% CI [1.25, 1.73], $p < 0.001$, std. $\beta$ = 4.60), with overall higher detection rates for person than plant stimuli. We also found a main effect of Age (exp. $\beta$ = 0.48, 95% CI [0.45, 0.52], $p < 0.001$, std. $\beta$ = -18.94) and a main effect of reported gender (exp. $\beta$ = 0.77, 95% CI [0.65, 0.91], $p < 0.001$, std. $\beta$ = -3.05). The interaction between object category and was also significant (exp. $\beta$ = 1.92, 95% CI [1.38, 2.66], $p < 0.001$, std. $\beta$ = 3.87). We then tested the inversion effect within each object category. The inversion effect was reliable for the detection rates of body stimuli ($\chi^2(1)$ = 29.61, $p < 0.001$, v = 0.07, OR = 2.00, 95 % CI = [1.56, 2.58], $BF_{10}$ > 1000). Upright bodies were detected by 97.2% of participants, while inverted bodies were detected by 94.5%. By contrast, the detection inversion effect was not reliably observed for plants (upright plant – 94.5%, inverted plant – 94.0 %; $\chi^2(1)$ = 0.66, $p$ = 0.42, v = 0.01, OR = 1.09, 95 % CI = [0.89, 1.34]), and the Bayesian analysis for this contrast indicated positive evidence for the null hypothesis ($BF_{10}$ = 0.04).

*Identification performance*

The results of identification performance are shown in Figure S3B. A logistic regression model on identification rates showed a main effect of Orientation (exp. $\beta$ = 4.37, 95% CI [3.97, 4.81], $p < 0.001$, std. $\beta$ = 30.01), of Object Category (exp. $\beta$ = 1.53, 95% CI [1.39, 1.69], $p < 0.001$, std. $\beta$ = 8.70), and of age (exp. $\beta$ = 0.77, 95% CI [0.73, 0.80], $p < 0.001$, std. $\beta$ = -12.72). Further, the Orientation x Category interaction was significant (exp. $\beta$ = 3.41, 95% CI [2.82, 4.14], $p < 0.001$, std. $\beta$ = 12.51). The effect of gender didn't reach significance ($p$ = 0.14). Chi-squared tests splitting the design by Category showed that upright person stimuli were identified correctly more often (93.3% of the times) than their inverted counterpart (64.0%; $\chi^2(1)$ = 852.93, $p < 0.001$, v = 0.36, OR = 7.85, 95% CI [6.73,

9.15], v = 0.25, $BF_{10} > 1000$). The inversion effect for the plant stimuli was smaller in size, but also reached significance (upright 83.3% vs inverted 68.0%, $\chi^2(1) = 216.35$, $p < 0.001$, OR = 2.34, 95 % CI [2.09, 2.63], v = 0.18, $BF_{10} > 1000$).

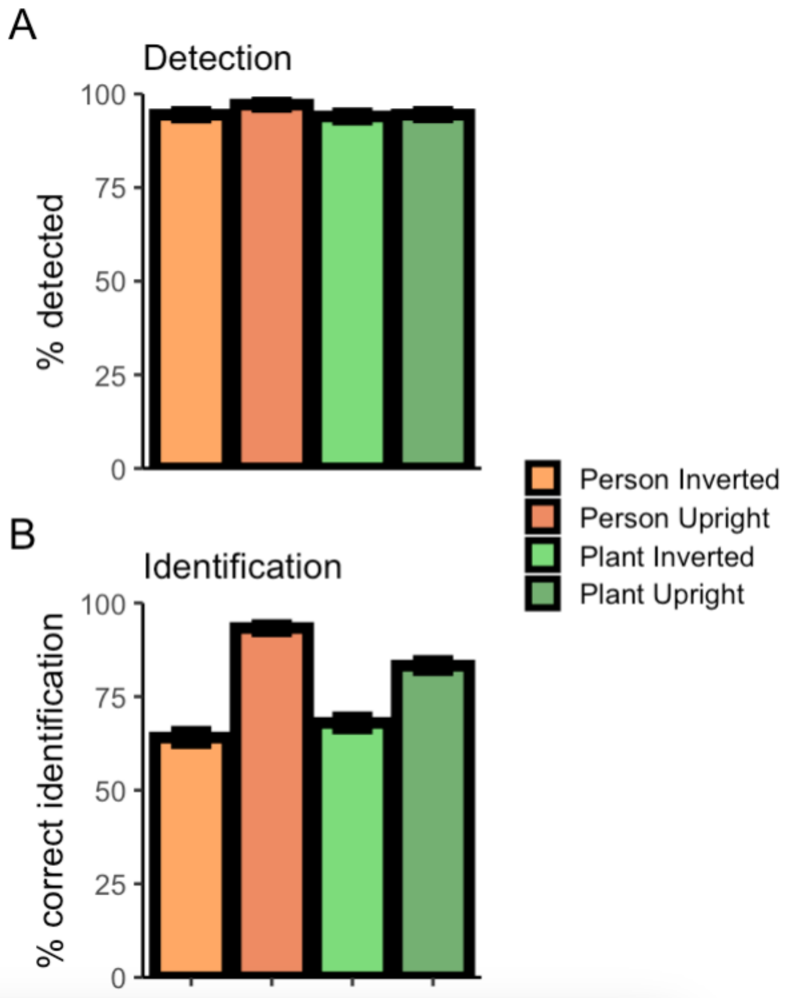


**Figure S3. A.** *Detection rates for Trial 8.* **B.** *Identification performance (% correct) for Trial 8.*

**Headless bodies – extended analyses**

We report here the exact same analysis approach as reported for the main experiment, now with headless bodies instead of full bodies.

**Methods**

Unless reported otherwise, methods were the same of the previous experiment.

**Participants**

The final sample considered for the analyses included N = 1412 (N = 853 identifying as woman, and N = 559 identifying as man; mean age = 34.2 ± 16.1; age range: 11-85 years). Details about participants' inclusion and exclusion criteria are provided in the Data analysis section. The data were collected between November 2024 and the beginning of February 2025. All participants gave consent for their data being used for scientific research. The procedures obtained approval from the Ethics Committee of the Faculty of Social Sciences at Radboud University (ECSW-2021-076).

**Stimuli**

The stimuli were 20 headless body silhouettes with the same luminance as those used in the main experiment (50%, middle gray RGB – 128, 128, 128). All bodies were standing, varied in their poses, and they were seen from different viewing angles. To add further stimulus variability, we also generated a horizontally flipped version of the stimuli. For each of these 40 headless body images we generated an inverted version by rotating the image by 180°. The plants were the same as in the previous experiment.

**Analyses**

*Participants exclusions.* The exclusions' criteria were the same as the main experiment. We considered population aged between 11-85 years old (N = 1825). We then excluded people who did not move the age slider from the default value of 50 (N = 112) and participants who declared to know already about the demonstration (N = 231). We then excluded participants who did not perform correctly the cross task in at least 2 out of 8 trials (N = 14) and participants who were 2.5 SDs slower than the group mean in the cross task (N = 16). To facilitate regressing out and to assess gender effects on a reasonably balanced sample we excluded datafiles in which the reported gender was "other" (i.e., neither man nor woman, N = 18) or that preferred to not report their gender (N = 22). Finally, for comparing the detection rates for each category and orientation with logistic regression we further excluded the participants who failed to detect a stimulus in Trial 8 (i.e. the full attention trial; N = 85).

**Results**

The difference between detection rates on the critical Trial 4 vs Trial 8 was significant ($\chi^2(1) = 675.57$, $p < 0.001$, v = 0.49, $BF_{10} > 1000$. In Trial 4, out of 1412 participants, 705 participants (49.9%) failed to detect the stimulus while being engaged in the cross task. Conversely, in Trial 8, the stimulus was detected by 1327 participants (94.0%) and only 6% of the participants (n = 85) failed to detect it. In Trial 7, more participants (1127 out of 1412; 79.8%) than in Trial 4 detected the presence of the stimulus ($\chi^2(1) = 274.4$, $p < 0.001$, v = 0.31, $BF_{10} > 1000$), meaning that even if they were not explicitly expecting another stimulus appearing besides the cross, the first detection question alone influenced detection rates. However, the detection rates in this trial were still reliably lower than in the control trial (Trial 8 - $\chi^2(1) = 124.41$, $p < 0.001$, v = 0.23, $BF_{10} > 1000$).

We then report performance in the cross task, to ensure that participants in this sample were also sufficiently engaged with the cross task. First, across trials, participants were able to perform the task above chance (M = 63.8%, SD = 18.3; t(1411) =28.26, $p < 0.001$), suggesting that they were sufficiently engaged with it. There was also an overall difference between the trials, as shown by a one-way ANOVA with trial number as a within-subject factor ($F(6.95, 9808.55) = 16.39$, $p < 0.001$, $\eta^2p = 0.1$). Similarly to the main experiment, the difference between performance in the cross task was numerically lower in Trial 7 (i.e., the divided attention trial, M = 65.23%, SD = 47.6) than in Trial 4, t(1411) = 1.93, $p = 0.054$, $BF_{10} = 0.19$), yet such difference did not reach significance. There was no difference when comparing Trial 4 (M = 68.63 %, SD = 46.4) and Trial 3 (M = 67.00%, SD = 47.0, t(1411) = 0.95, $p = 0.34$, $BF_{10} = 0.05$), confirming that participants continued to prioritize the cross task on Trial 4.

***Trial 4 – No expectation***

*Detection rates*

Here we tested the same logistic models as used in the main experiment. A logistic regression model on detection rates showed a main effect of orientation (exp. $\beta = 1.28$, 95% CI [1.02, 1.61], $p = 0.030$, std. $\beta = 2.17$), with higher detection rates for upright than inverted stimuli. We also found a main effect of Age (exp. $\beta = 0.58$, 95% CI [0.52, 0.66], $p < 0.001$, std. $\beta = -8.97$) and a main effect of reported gender (exp. $\beta = 0.65$, 95% CI [0.51, 0.82], $p <$ 0.001, std. $\beta = -3.66$). The interaction between object category and orientation approached, but did not reach significance, likely due to the much smaller sample size of this additional experiment compared to the main experiment (exp. $\beta = 1.48$, 95% CI [0.94, 2.33], $p = 0.087$, std. $\beta = 1.71$). Due to the between-subjects nature of the design we tested the inversion effect within each object category. The inversion effect was reliable for the detection rates of body stimuli ($\chi^2(1) = 7.11$, $p = 0.008$, v = 0.11, OR = 1.54, 95 % CI = [1.13, 2.09], $BF_{10} =$ 8.30). Upright bodies were detected by 59.1% of participants, while inverted bodies were detected by 48.5%. By contrast, the detection inversion effect was not reliably observed for plants (upright plant – 50.5%, inverted plant – 49.1 %; $\chi^2(1) = 0.09$, $p = 0.77$, v = 0.01, OR = 1.06, 95 % CI = [0.78, 1.44]), and the Bayesian analysis for this contrast indicated positive evidence for the null hypothesis ($BF_{10} = 0.21$).

*Identification performance*

A logistic regression model on identification performance showed a main effect of Orientation (exp. $\beta = 1.94$, 95% CI [1.55, 2.42], $p < 0.001$, std. $\beta = 5.79$). Further, the Orientation x Category interaction was significant (exp. $\beta = 2.08$, 95% CI [1.33, 3.25], $p =$ 0.001, std. $\beta = 3.21$). No other effect reached significance (all *ps* > 0.10). Chi-squared tests splitting the design by Category showed that upright bodies were identified correctly more often (68.66% of the times) than their inverted counterpart (43.60%; $\chi^2(1) = 41.28$, $p < 0.001$, v = 0.30, OR = 2.83, 95% CI [2.06, 3.89], v = 0.25, $BF_{10} > 1000$). The inversion effect for the plant stimuli was smaller and approached, but did not reach significance (upright 62.89% vs inverted 55.50%, $\chi^2(1) = 3.39$, $p = 0.066$, OR = 1.36, 95 % CI [0.99, 1.86], v = 0.07, $BF_{10} =$ 1.19).

***Trial 7 – Some expectation***

*Detection rates*

A logistic regression model on detection rates in Trial 7 showed a main effect of orientation (exp. $\beta = 1.65$, 95% CI [1.23, 2.23], $p < 0.001$, std. $\beta = 2.17$), with higher detection rates for upright than inverted stimuli. We also found a main effect of Age (exp. $\beta = 0.57$, 95% CI [0.50, 0.66], $p < 0.001$, std. $\beta = -7.78$). The interaction between object category and orientation did not reach significance, likely due to the much smaller sample size of this additional experiment compared to the main experiment (exp. β = 1.52, 95% CI [0.84, 2.76], *p* = 0.167, std. β = 1.38). Due to the between-subjects nature of the design we tested the inversion effect within each object category. The inversion effect was reliable for the detection rates of body stimuli ($\chi^2(1) = 6.31$, *p* = 0.012, v = 0.11, OR = 1.79, 95 % CI = [1.15, 2.78], $BF_{10} = 4.51$). Upright bodies were detected by 88% of participants, while inverted bodies were detected by 80.4%. By contrast, the detection inversion effect was not reliably observed for plants (upright plant – 82.8%, inverted plant – 78.3 %; $\chi^2(1) = 1.98$, *p* = 0.160, v = 0.10, OR = 1.34, 95 % CI = [0.92, 1.95], $BF_{10} = 0.46$).

*Identification performance*

A logistic regression model on identification performance showed a main effect of Orientation (exp. β = 3.95, 95% CI [3.07, 5.10], *p* < 0.001, std. β = 10.58). The main effect of age was also significant (exp. β = 0.85, 95% CI [0.75, 0.96], *p* = 0.007, std. β = -2.70). Further, the Orientation x Category interaction was significant (exp. β = 5.47, 95% CI [3.29, 9.10], *p* < 0.001, std. β = 6.55). No other effect reached significance (all *ps* > 0.23). Chi-squared tests splitting the design by Category showed that upright bodies were identified correctly more often (86.69% of the times) than their inverted counterpart (42.33%; $\chi^2(1) = 41.28$, *p* < 0.001, v = 0.30, OR = 2.83, 95% CI [2.06, 3.89], v = 0.25, $BF_{10} > 1000$). The inversion effect for the plant stimuli was smaller but significant (upright 70.91% vs inverted 59.04%, $\chi^2(1) = 10.76$, *p* = 0.001, OR = 1.69, 95 % CI [1.23, 2.32], v = 0.12, $BF_{10} = 38.75$).

***The role of age and reported gender***

*Detection rates*

We analyzed in this additional and smaller subset of participants whether we could reproduce the same pattern of effects of age and reported gender on detection rates in inattentional blindness. The best model explaining detection rates (averaged across object category and orientation of the stimulus) was the same as the one in the main experiment and included the three main effects of Age, Gender, and Trial. Including the interaction between Age and Gender, or between all three conditions, did not further improve the model fit (all *p*s from the likelihood ratio tests comparing the models with two- or three-way interactions to the main effect model were *p* > 0.259). For both trials (Trial 4 – M = 57.3%, W = 48.1%, Trial 7 – M = 84.3, W = 81.0), people identifying themselves as men were 1.31 times more likely to detect a stimulus than women (exp. β = 1.31, 95% CI [1.17, 1.42], *p* < 0.001, std. β = 3.93). Again, this gender difference was present despite the fact that performance of the two groups did not differ in the main cross task – (between subjects tests by gender in Trial 4 – *t*(1325) = -0.73, *p* = 0.467, $BF_{10} = 0.08$, Trial 7 – t(1325) = 0.71, *p* = 0.479, $BF_{10} = 0.08$, and on overall performance, t(1325) = -0.68, *p* = 0.498, $BF_{10} = 0.03$), suggesting that the increased inattentional blindness in women is not explained by gender differences in the primary cross task performance.

Concerning the age differences, the older were the participants the less likely they were to detect the stimulus appearing in the cross (exp. β = 0.58, 95% CI [0.53, 0.64], *p* < 0.001, std. β = -11.80). To facilitate visualization of the age differences, we plotted the model's predicted probabilities of detecting the stimulus for both Trial 4 and Trial 7 (see **Figure S4**). The probability of noticing a stimulus in the absence of expectation decreases on average by 7.33 % (SD = 1.31) every 10 years of increased age. In the presence of some expectation (Trial 7), we observe an average decrease of 6.28% in detection every 10 years (SD = 1.99). There was, however, a small but reliable negative correlation between age and performance in the cross task ($\rho$ = -0.06, $p$ = 0.02, $BF_{10}$ = 5.21).

*Identification performance*

The same model on identification performance showed a main effect of Age (exp. $\beta$ = 0.90, 95% CI [0.83, 0.98], $p$ = 0.01, std. $\beta$ = -2.59). The probability of identifying the stimulus in absence of expectation (Trial 4) decreased on average by 1.61 % (SD = 0.03) every 10 years of increased age, while in Trial 7 by 1.52% (SD = 0.06). Differently from the detection rates, the participants' gender did not reliably predict identification performance (exp. $\beta$ = 0.95, 95% CI [0.77, 1.11], $p$ = 0.563, std. $\beta$ = -0.58).

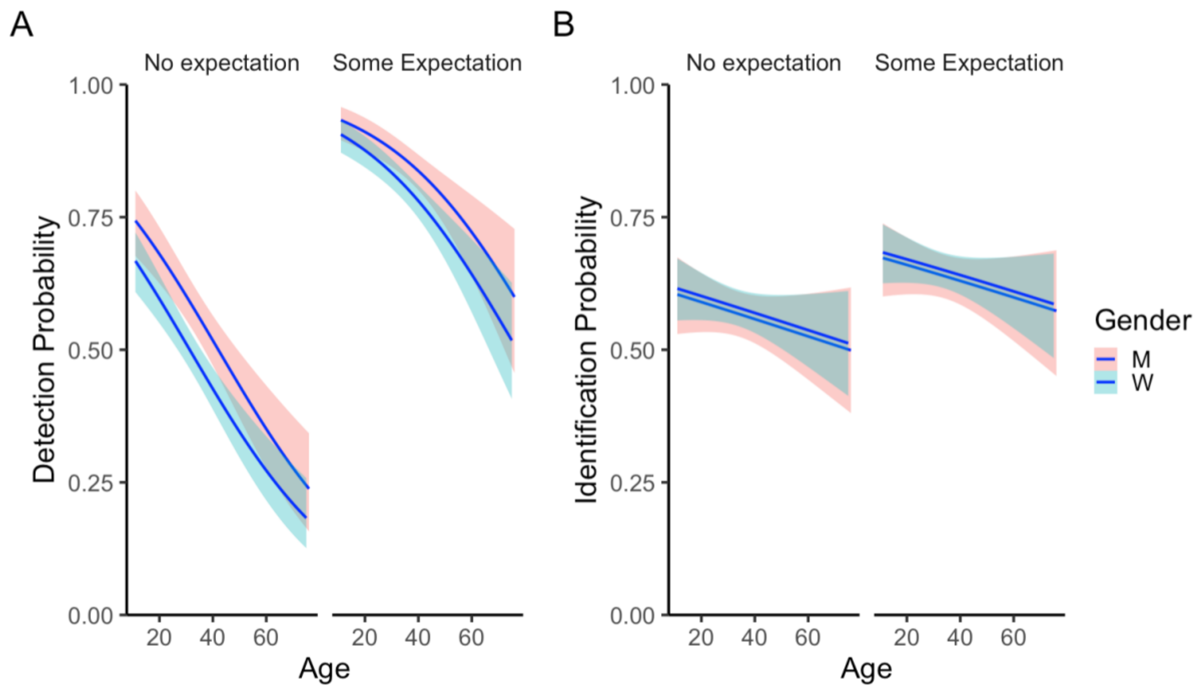


**Figure S4. A.** *Estimated probability of detection based on the participants' reported age and gender. The left panel shows the detection probability by age split between people identifying as women (light blue) and men (light red) in Trial 4 (No expectation). The right panel shows the estimate for Trial 7 (Some expectation).* ***B.*** *Estimated probability of identification based on the participants' reported age and gender. The left panel shows the detection probability by age split between people identifying as women and men in Trial 4 (No expectation). The right panel shows the estimate for Trial 7 (Some expectation). The shaded area indicates the 95% confidence interval of the estimate.*

#### *Comparison between bodies with and without heads*

When running logistic regression models to predict detection rates by Stimulus Category (Body with Heads/Headless bodies) x Orientation (Upright/Inverted) we consistently found a main effect of orientation (Trial 4 detection: exp. $\beta$ = 1.44, 95% CI [1.22, 1.71], $p$ < 0.001, std. $\beta$ = 4.31; Trial 7 detection: exp. $\beta$ = 1.89, 95% CI [1.49, 2.39], $p$ < 0.001, std. $\beta$ = 5.26; Trial 4 Identification: exp. $\beta$ = 3.09, 95% CI [2.61, 3.65], $p$ < 0.001, std. $\beta$ = 13.09; Trial 7

Identification: exp. β = 7.98, 95% CI [6.43, 9.91], $p < 0.001$, std. β = 18.86), and not reliable interaction between Object Category (Person with head/headless) and Orientation(Trial 4 detection: exp. β = 0.88, 95% CI [0.63, 1.23], $p = 0.442$, std. β = -0.77; Trial 7 detection: exp. β = 0.85, 95% CI [0.53, 1.36], $p = 0.493$, std. β = -0.69; Trial 4 identification: exp. β = 1.26, 95% CI [0.90, 1.78], $p = 0.185$, std. β = 1.33; Trial 7 identification: exp. β = 0.68, 95% CI [0.45, 1.04], $p = 0.072$, std. β = -1.80), suggesting that the inversion effects of bodies with and without heads were comparable in size.

**Identification performance on detected stimuli**

When looking at the identification performance only for participants who successfully detected the stimuli, we observed the same pattern as when analyzing all participants (including those who did not detect the stimuli). That is, stimulus inversion affected identification performance of both the person and the plants, and to a larger extent affected the recognition of person stimuli. In both Trial 4 and Trial 7, there was the same stimulus x orientation interaction (all *p*s from the logistic models were < 0.001). As one would expect, when selecting only those participants that correctly detected the stimulus, the effect of inversion for both the bodies and the plants became even larger in percentage than when considering all of the participants (difference between upright and inverted Person in trial 4: 40.6 % – instead of 28.0 %; Plant in trial 4: 13.72% - instead of 6.63%; Person in trial 7: 35.7% - instead of 32.79%; Plant in trial 7: 16.1 % - instead of 14.2 %).

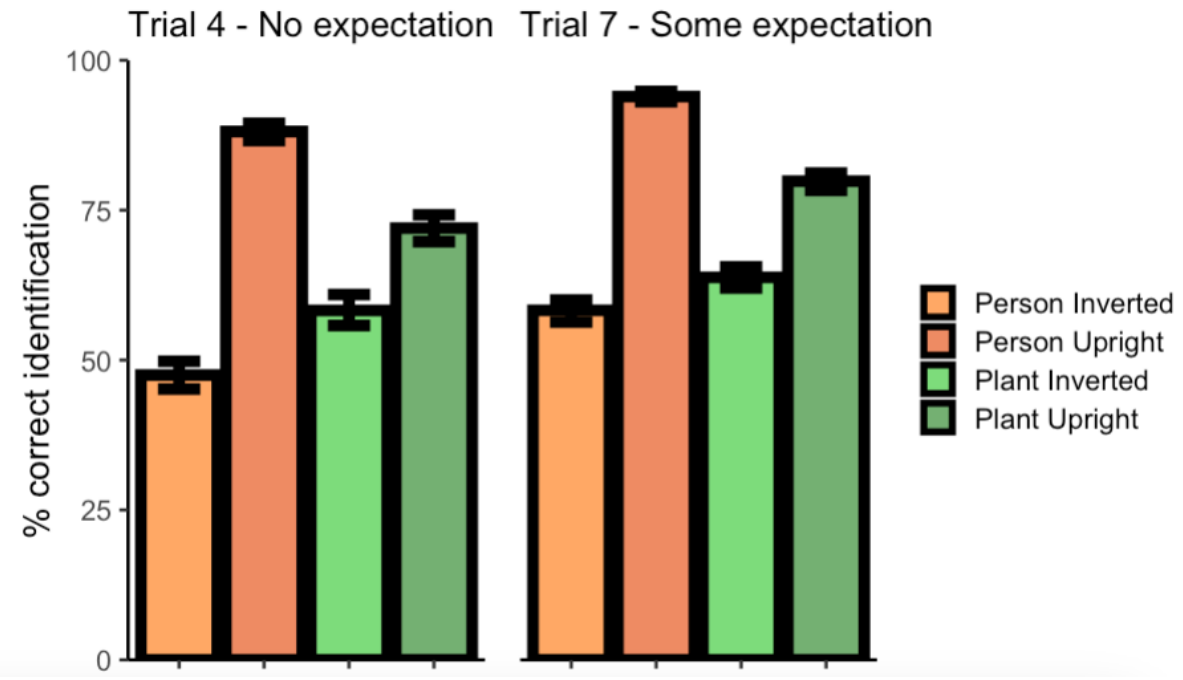


**Figure S5 –** *Identification performance for Trial 4 (left panel) and Trial 7 (right panel) for participants who had detected the stimuli in that trial*.

**Plots of Detection in Trial 7 as a function of the observed stimulus in Trial 4**

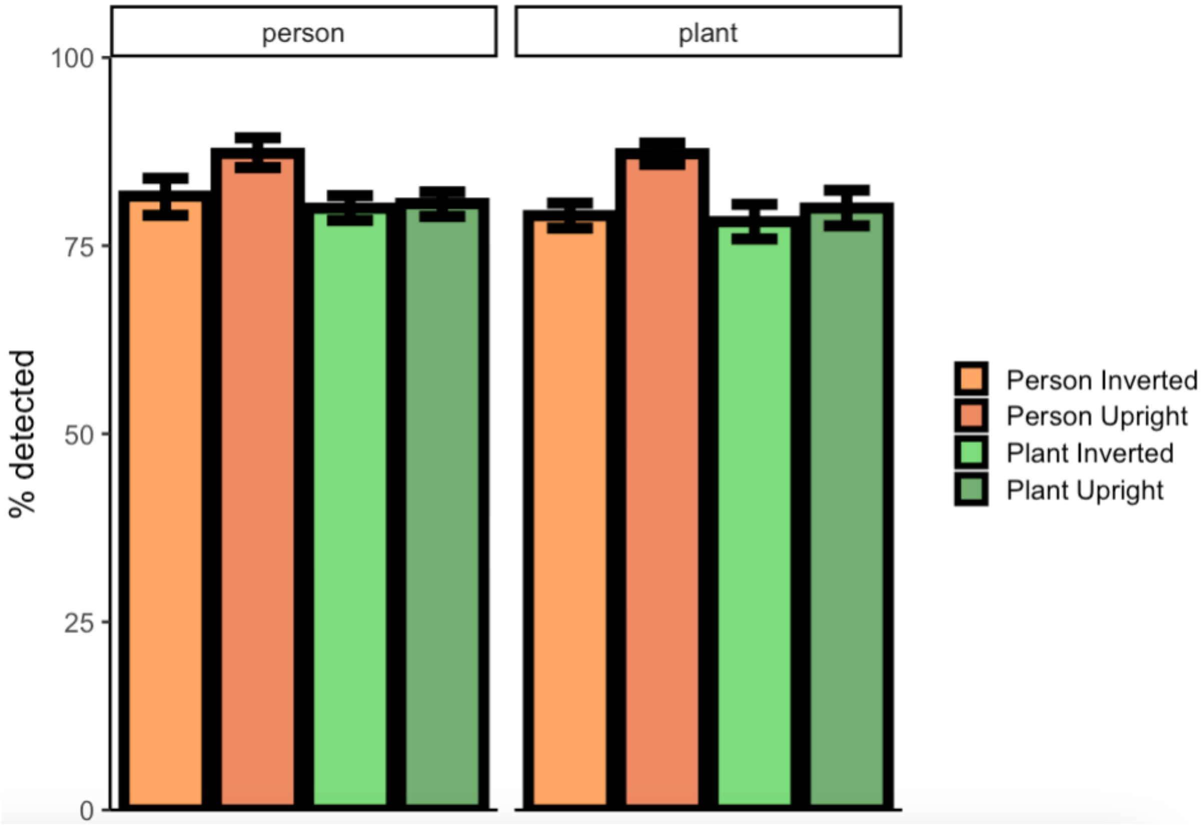


**Figure S6 –** *Detection rates in Trial 7 as a function of the stimulus that appeared in Trial 4 (left panel: a person appeared in Trial 4; right panel: a plant appeared in Trial 4).*

### Supplementary references


Akaike, H. (1981). Likelihood of a model and information criteria. *Journal of econometrics*, *16*(1), 3-14.

Bozdogan, H. (1987). Model selection and Akaike's information criterion (AIC): The general theory and its analytical extensions. *Psychometrika*, *52*(3), 345-370.

Darlington, R. B. (1990). *Regression and linear models*. McGraw-Hill Companies.

Hosmer Jr, D. W., Lemeshow, S., & Sturdivant, R. X. (2013). *Applied logistic regression*. John Wiley & Sons.

Stoltzfus, J. C. (2011). Logistic regression: a brief primer. *Academic emergency medicine*, *18*(10), 1099-1104.
